# A more accurate account of the effect of k-space sampling and signal decay on the effective spatial resolution in functional MRI

**DOI:** 10.1101/097154

**Authors:** Denis Chaimow, Amir Shmuel

## Abstract

The effects of k-space sampling and signal decay on the effective spatial resolution of MRI and functional MRI (fMRI) are commonly assessed by means of the magnitude point-spread function (PSF), defined as the absolute values (magnitudes) of the complex MR imaging PSF. It is commonly assumed that this magnitude PSF signifies blurring, which can be quantified by its full-width at half-maximum (FWHM). Here we show that the magnitude PSF fails to accurately represent the true effects of k-space sampling and signal decay.

Firstly, a substantial part of the width of the magnitude PSF is due to MRI sampling per se. This part is independent of any signal decay and its effect depends on the spatial frequency composition of the imaged object. Therefore, it cannot always be expected to introduce blurring. Secondly, MRI reconstruction is typically followed by taking the absolute values (magnitude image) of the reconstructed complex image. This introduces a non-linear stage into the process of image formation. The *complex* imaging PSF does not fully describe this process, since it does not reflect the stage of taking the magnitude image. Its corresponding *magnitude* PSF fails to correctly describe this process, since convolving the original pattern with the magnitude PSF is different from the true process of taking the absolute following a convolution with the complex imaging PSF. Lastly, signal decay can have not only a blurring, but also a high-pass filtering effect. This cannot be reflected by the strictly positive width of the magnitude PSF.

As an alternative, we propose to first approximate the MRI process linearly. We then model the linear approximation by decomposing it into a signal decay-independent MR sampling part and an approximation of the signal decay effect. We approximate the latter as a convolution with a Gaussian PSF or, if the effect is that of high-pass filtering, as reversing the effect of a convolution with a Gaussian PSF. We show that for typical high-resolution fMRI at 7 Tesla, signal decay in Spin-Echo has a moderate blurring effect (FWHM = 0.89 voxels, corresponds to 0.44 mm for 0.5 mm wide voxels). In contrast, Gradient-Echo acts as a moderate high-pass filter that can be interpreted as reversing a Gaussian blurring with FWHM = 0.59 voxels (0.30 mm for 0.5 mm wide voxels). Our improved approximations and findings hold not only for Gradient-Echo and Spin-Echo fMRI but also for GRASE and VASO fMRI. Our findings support the correct planning, interpretation, and modeling of high-resolution fMRI.

## Introduction

The spatial specificity of functional MRI (fMRI) based on the Blood Oxygenation Level Dependent (BOLD) signal depends on the spatial properties of the hemodynamic response. Specifically, it depends on the relative contributions of the micro-vascular and macro-vascular components of the hemodynamic response to the fMRI signal. In addition to the effects of the hemodynamic response on the spatial specificity of fMRI, the MRI acquisition process influences the effective resolution of the acquired images. Specifically, the sampling of k-space by means of temporal gradient encoding defines the spatial resolution. However, the effective spatial resolution can be compromised in the presence of *T*_2_ and/or 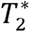 decay, which potentially contribute to the overall measured spread of the BOLD fMRI signal.

The BOLD point-spread function (PSF) is a measure used to approximate the spatial spread of the BOLD response to a localized increase in neuronal activity. A convolution of the pattern of neuronal activity with a single BOLD PSF kernel is not a precise model of the spatial specificity of the BOLD response, because of the variability in the vascular components as a function of space (Polimeni et al., 2010). However, it provides a useful measure, based on the average BOLD PSF across space, for comparing the spatial specificity between different fMRI contrasts and techniques.

The full-width at half-maximum (FWHM) of the gradient echo (GE) BOLD PSF at 1.5 T was found to be 3.5 mm (Engel et al., 1997). Similar values of 3.9 mm for GE BOLD and 3.4 mm for Spin-Echo (SE) BOLD have been reported at 3 T (Parkes et al., 2005). We previously estimated the FWHM of the GE BOLD PSF to be below 2 mm at 7 T (Shmuel et al., 2007). Narrower BOLD PSFs at higher field strength are thought to result from reduced intravascular contributions from larger blood vessels and increases in extravascular signal changes around capillaries and smaller vessels (Yacoub et al., 2001). Additional relative weighting towards the microvasculature, and thus further increases in spatial specificity, can be achieved by using SE BOLD imaging, which suppresses extravascular signal contributions from larger blood vessels (Uludağ et al., 2009; Yacoub et al., 2003).

The use of high field strengths and developments in pulse sequences that lead to decreases in the BOLD fMRI PSF allow investigation into the function of ever finer structures such as cortical columns (Cheng et al., 2001; Goodyear and Menon, 2001; Menon et al., 1997; Nasr et al., 2016; Shmuel et al., 2010; Yacoub et al., 2008; 2007; Zimmermann et al., 2011;). Consequently, in order to optimize such experiments and to understand their inherent limitations, it becomes important to assess the contribution of the MRI sampling process and of 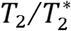 decay to the overall BOLD fMRI PSF.

The MR imaging process can be described by means of a complex valued MR imaging PSF (Haacke et al., 1999). The magnitude PSF (formed by the absolute values (magnitudes) of the complex MR imaging PSF) and the corresponding FWHM of the magnitude PSF, have been used to assess the effective spatial resolution and to quantify the blurring that the MR sampling process introduces (Constable and Gore, 1992; Farzaneh et al., 1990; Haacke, 1987; Kemper et al., 2015; Oshio and Singh, 1989; Qin, 2012).

Here, we simulate 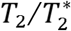 decay and MR imaging of realistic columnar patterns to show that the FWHM of the magnitude PSF is neither a meaningful nor an accurate measure for quantifying the effect of MR sampling on the effective spatial resolution, especially in the context of functional MRI. As an alternative, we propose to decompose the modeling of the imaging process into two components: one component accounts for MR sampling, independent of the signal decay; a second component, formulated as a convolution with a Gaussian kernel, approximates the blurring effect due to the 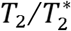 decay.

## Methods

### Discrete representations of simulated spaces

All simulations were implemented in MATLAB (The MathWorks Inc., Natick, MA, USA). Spatial dimensions were considered relative to an arbitrary voxel width. A field-of-view (FOV) of 32 voxels was simulated, represented by 256 equally spaced points (resolution 8 times finer than the voxel width). The spatial frequency space (k-space) was simulated on a corresponding grid of 256 equally spaced points representing a spatial frequency range between -128 and +127 cycles per 32 voxels. Spatial frequencies sampled by MRI (see below) are represented by the central part of this simulated k-space, covering the spatial frequency range between -16 and +15 cycles per 32 voxels.

### Modeling of signal decay

GE signal decay *f*_*GE*_(*t*) and SE signal decay *f*_*SE*_(*t*) were modeled according to (Haacke et al., 1999): 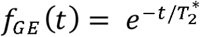

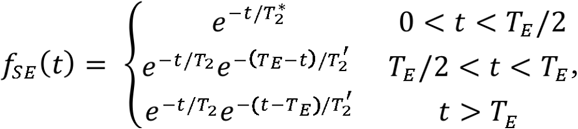

where *T*_*E*_ represents the echo time. Relaxation time constants for gray matter at 7T were used (see Uludağ et al., 2009 for a review of relaxation times). *T*_2_ was set to 50 ms. 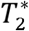 of gray matter at 7T was 27.8 ms. In order to account for additional macroscopic inhomogeneities, a volumetric 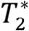 value of 17 ms was used (Kemper et al., 2015).

### Calculation of Modulation Transfer Functions

If not stated otherwise, a total read-out time of 27.8 ms (equal to 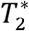) for the full k-space acquisition using 32 lines (phase-encode steps) was assumed. For partial Fourier acquisition, the total read-out time was shortened, so that it was proportional to the reduction in k-space coverage, resulting in a total read-out time of 20.85 ms for the acquisition of 24 lines (3/4 partial Fourier).

For GE, the echo time *T*_*E*_ was set to the true 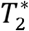 of 27.8 ms. The SE echo time was set to 55 ms (Yacoub et al., 2003). The modulation transfer functions (MTFs) of MR imaging are sums of Dirac delta functions (rect-function-windowed Dirac comb function), where each Dirac delta function is modulated by a signal decay factor (Appendix A). Discrete representations of MTFs were computed by first calculating the sampling time of each k-space line relative to excitation and then setting the line s MTF value to the signal decay value for this time, or to zero if it fell outside the range of sampled lines.

### Calculation of point-spread functions

PSFs were calculated by taking an inverse discrete Fourier transform of the discrete representation of the corresponding MTF.

### Simulation of responses of cortical columns

MR imaging simulations were applied to simulated realistic ocular dominance column (ODC) patterns and to simulated general isotropic columnar patterns.

ODC response patterns were simulated by anisotropic filtering of Gaussian white noise (Rojer and Schwartz, 1990). Detailed modeling equations can be found in (Chaimow et al., 2011). We modified the mathematical form of the band-pass filtering kernel relative to our previously published model, expressing it as a product of radial and angular components. The non-normalized filter as a function of absolute spatial frequency and orientation ϕ is given as

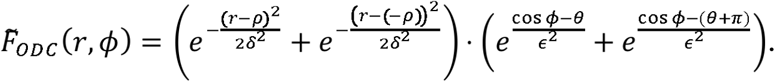

Unless stated otherwise, the main spatial frequency parameter ρ was set to 0.5 cycles/mm, corresponding to an average column width of 1 mm (Yacoub et al., 2007). Parameters and controlled the degree of irregularities orthogonal and parallel to the main axis of elongation of the ODC bands, respectively. We specified those parameters by defining *relative irregularity* parameters as 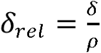 and 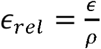, that determined the level of irregularities independent of the chosen main spatial frequency. Unless stated otherwise, relative irregularity parameters σ_ret_ and ε_ret_ were set to 0.5 and 1 respectively. The orientation parameter θ was set to π/2, so that the main axis of elongation of the ODC bands was orthogonal to the phase-encode direction.

For 1D modeling (along the phase-encode direction), the 2D model was reduced by only considering the radial component of the filter (first factor), setting the angular component (second factor) to 1. This 1D model can be regarded as a general one dimensional columnar model, valid not only for anisotropic organizations such as ODC, but also for isotropic columnar patterns.

The sharpness parameter α was set to 1.4, resulting in a mid-level of sharpness. The default maximum response amplitude was set to 5%. Single condition columnar response patterns were added to a background signal intensity of 1. For the definition of the sharpness parameter and the response amplitude, see (Chaimow et al., 2011).

### Simulation of MR imaging

MR imaging was modeled by multiplying the k-space representation of a pattern with the MTF and applying an inverse discrete Fourier transformation. This was followed by taking the magnitude of the resulting complex values. The size of the MTF was identical to that of the pattern k-space representation. However, the MTF was zero for k-space components not sampled by MRI imaging. Therefore the result of the discrete Fourier transform was a high-resolution representation of the voxel-size dependent MR sampled signal. It is equivalent to an interpolation using zero-filling in k-space. This high-resolution representation was then down-sampled to correspond to the actual voxel size, resulting in single voxel signals in accordance with the MR imaging equation (Appendix A).

Different MTFs introduce differences in the overall amplitude scaling of resulting images. To allow the comparison of the image patterns obtained by considering different MTFs, we normalized the magnitude images in the last stage of the simulation. To this end, we divided each of the magnitude images by a constant equal to the result of simulating the entire MRI sampling process with the specific MTF applied to a constant pattern of value 1. Note that 1 is also the value of the homogeneous background onto which we superimposed the simulated ODC pattern with the maximal amplitude of 5%.

### Simulation of Partial Fourier

Partial Fourier imaging and reconstruction was simulated by setting the first or last ¼ of the MTF components within the full acquisition sampling range to either zero (zero-filling reconstruction) or to their conjugate symmetric counterparts (conjugate symmetry reconstruction). The most negative k-value of the full acquisition sampling range has no positive k-space counterpart due to the slight asymmetric sampling of an even number of k-space lines. Therefore, in early omission partial Fourier and conjugate symmetry reconstruction, the most negative k-value was set to zero.

### Implementation of approximating MRI models

Convolutions between simulated patterns and various kernel functions were implemented as multiplication of their respective discrete Fourier transforms or MTF representations, followed by inverse discrete Fourier transform back into image space. For Gaussian blurring, Gaussian kernels were computed as *e*^-*x*^2^/2*σ*^2^^ where *σ* = *FWHM*/2.355.

Convolutions followed by MR sampling were modeled by first setting the MTF outside the range of the sampled lines to zero. Then, this modified MTF was multiplied with the k-space representation of the pattern, and an inverse discrete Fourier transformation was applied. Lastly, the magnitudes of the resulting complex values were computed. This high-resolution representation was then down-sampled according to the voxel size, resulting in single voxel signals according to the MR imaging equation (Appendix A).

### Contrast range

The contrast range of each of 1000 simulated individual response patterns, superimposed on a background signal intensity level of 1, was computed as the standard deviation (SD) of all responses while taking into account the theoretical mean(1 + *β*2, where *β* is the maximum response level) used for simulating the original pattern. All individual contrast range estimates were averaged resulting in the average contrast range.

### Frequency spectra

Spatial frequency spectra were computed by taking the absolute value of the discrete Fourier transform separately for each of 1,000 individual response patterns. Average spatial frequency spectra were computed by averaging all individual spectra.

### Evaluation of a linear approximation of the MR imaging process

We approximated the MR imaging process linearly as a convolution with the real component of the complex imaging PSF (see Appendix B). This linear approximation of the MR imaging process was evaluated by simulating 1000 different cortical columnar response patterns for each combination of a main spatial frequency (8 values from 1 cycle per Field of View (FOV) to 1 cycle per 2 voxels), a relative irregularity parameter (10 values from 0.1 to 1), and a range of maximum response amplitudes (1% -10% in steps of 1% and 10% -100% in steps of 10%). The complete MR imaging of each of these patterns was simulated. In addition, for each of these patterns we computed the linear approximation of the MR imaging process. The results of the linear approximations of the MR imaging process were compared to the complete MR imaging simulations by computing the root-mean-squared-errors (RMSE) relative to the standard deviation of the simulated patterns of the complete MR imaging process.

In addition, for a response amplitude of 5%, the same patterns were also convolved with the magnitude PSF. The results of convolving with the magnitude PSF were compared to the complete MR imaging simulations by computing the root-mean-squared-errors (RMSE) relative to the standard deviation of the simulated patterns of the complete MR imaging process.

### Definition and fitting of a Gaussian point-spread function model for signal decay

The MTF corresponding to the real component of the complex imaging PSF 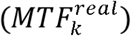 was calculated by transforming the real component of the complex imaging PSF back into the spatial frequency domain. The inverse MTF of this real component 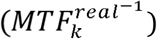 was calculated as 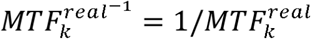 for all k-space lines within the sampling range (and zero outside the sampling range).

Gaussian functions of the form 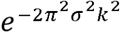 were fitted to 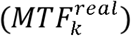 and 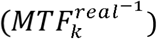 within the range of –(k-1) to (k-1). The amplitude *a* was constrained to be equal to the center component 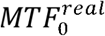 or 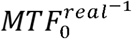. This resulted in Gaussian PSFs of the form 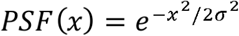 whose FWHM = *σ* 2.355. To fit the Gaussian functions, we used the MATLAB function ‘fit from the Curve Fitting Toolbox (The MathWorks Inc., Natick, MA, USA). The Gaussian fit (either to the real component of the complex imaging PSF or to its inverse) with the higher R^2^ was considered for further characterizing the effects of signal decay.

### Evaluation of the Gaussian point-spread function model of signal decay

The Gaussian PSF model of signal decay was evaluated by simulating 1000 different cortical columnar response patterns with a main spatial frequency of 1 cycle per 4 voxels and a relative irregularity of 0.5. The complete MR imaging of these patterns was simulated for different total read-out durations and partial Fourier acquisition schemes (including full k-space acquisition).

In addition, for each of these patterns and acquisition parameters (different total read-out durations and partial Fourier acquisition schemes), we computed the convolution of the pattern with a Gaussian PSF model for the signal decay (see definition and fitting described in the previous section). This was followed by MR sampling with no decay. The results of these approximations of the MR imaging process were compared to the complete MR imaging simulations by computing the root-mean-squared-errors (RMSE) relative to the standard deviation of the simulated patterns of the complete MR imaging process.

### Estimation of a pattern specific Gaussian point-spread function model of signal decay

One thousand different cortical columnar response patterns with a main spatial frequency of 1 cycle per 4 voxels and a relative irregularity of 0.5 were simulated. MR imaging of these patterns was simulated for different total-read out durations and partial Fourier acquisition schemes (including full k-space acquisition). For each pattern, we computed convolutions with PSFs corresponding to Gaussian MTFs and inverse of Gaussian MTFs, while considering the FWHM as a free parameter. This was followed by MR sampling with no decay. Using MATLAB s fminsearch (The MathWorks Inc., Natick, MA, USA), the FWHM (and the choice of Gaussian MTF or inverse of Gaussian MTF) was optimized such that the mean-squared-error between the approximation and the full MR imaging simulation was minimized.

## Results

### The MR imaging point-spread function

We first summarize how the MRI acquisition process of a pattern can be described using PSFs. Appendix A provides detailed equations. The theory follows Haacke et al. (1999).

### MRI with no signal decay

Echo-planar imaging (EPI) samples the two-dimensional k-space representation of the pattern by sequentially sampling individual lines along the first dimension (read-out direction), each separated by a step in the phase-encode direction. This results in a grid of sampled k-space points from which the original image is reconstructed using an inverse discrete Fourier transform. Since the dimensions in the Fourier transform are separable, we can focus on one dimension: the phase-encode dimension.

Assuming no signal decay takes place, we can formulate the MRI sampling process as an inverse Fourier transform of the product between the k-space representation of the pattern and a rect-function-windowed Dirac comb function (Figure 1, no decay, MTF). This function describes the effect of a linear system as multiplication in spatial frequency space and is commonly termed a modulation transfer function (MTF). Typically an even number of points, *N* = 2*n* is sampled, resulting in a slightly asymmetric coverage of k-space over the region [-*n*Δ*k*, (*n* - 1) Δ*k*], where Δ*k* is the step size in the k-space.

**Fig. 1.**
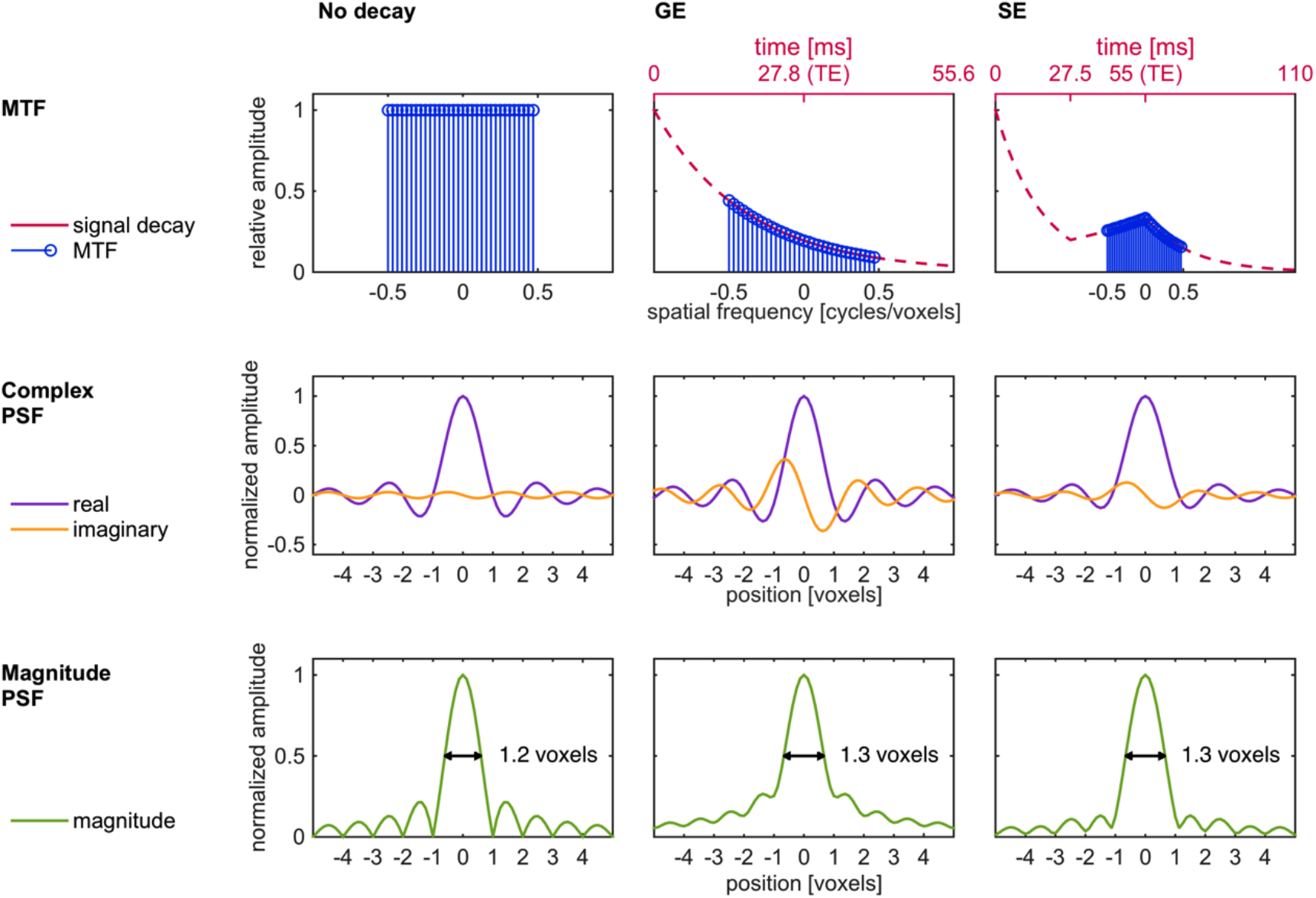
MR complex imaging PSF and its absolute values. This figure demonstrates how MR imaging can be described using a PSF. The three columns illustrate the scenarios of no signal decay, signal decay in GE imaging, and signal decay in SE imaging, respectively. The first row shows the modulation transfer function (MTF) of the imaging process. In the case of no decay, the MTF is a Dirac comb function that corresponds to sampled k-space data points that are sampled in time from the lowest to the highest k-value. In GE imaging, this comb function is modulated by the 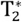 signal decay (red line). In SE a refocusing pulse results in a decay curve with a peak at the echo time. The second row shows the corresponding complex PSFs defined as the Fourier transform of the MTF. These functions show the complex signal one would obtain along the phase encoding direction from an infinitesimally small point-like structure. One can model the complex signal obtained by MRI along the phase-encoding direction as a convolution of the original pattern with the complex PSF. In contrast, in the general case one cannot model the magnitude of the signal obtained along the phase-encoding direction as the result of a convolution process. The third row shows the absolute (magnitude) values of the complex imaging PSF. Its FWHM (black arrows) has been previously used as a common measure to describe the spatial specificity of the MR imaging process. For relaxation time constants measured at 7 Tesla and a total readout duration of 27.8 ms, the FWHM of the magnitude PSF is 1.20 voxels for no decay, 1.34 voxels for GE, and 1.32 voxels for SE.

Multiplying the k-space data with an MTF is equivalent to convolving the image space data with the MTF s Fourier transform, which is the imaging point-spread function (PSF) (Figure 1, no decay, complex PSF). On its own, the PSF describes the image one would obtain from an infinitesimally small point-like structure.

The non-zero imaginary component of the no-decay PSF is a result of the above mentioned asymmetry in the MTF. MTF asymmetries are also caused by signal decay as described below. MTF asymmetries and other phase-influencing artifacts result in reconstructed images that are generally complex valued with non-zero phase. Commonly, the absolute values (magnitudes) of the complex image are considered for further analysis. Likewise, the spatial resolution of MRI and fMRI is often characterized by measuring the full-width at half-maximum of the magnitude PSF, obtained by taking the absolute values (magnitudes) of the complex PSF (Figure 1, no decay, magnitude PSF). Here, without any signal decay effects the FWHM is 1.20 voxels.

However, neither the complex PSF nor the magnitude PSF can describe the spatial resolution of the MRI process correctly under all circumstances. The magnitude PSF in itself is not the PSF of the imaging process, because convolution with the absolute values of the complex PSF is not equivalent to taking the absolute values after convolution with the complex PSF (which is the common practice in MRI reconstruction). In contrast, the complex PSF does describe the imaging process (excluding the operation of taking the absolute values of the complex image). However, given its complex nature, how to use the complex PSF for quantifying the effective spatial resolution of the absolute values image is not obvious.

### Signal decay in Gradient-Echo and Spin-Echo functional MRI

So far, we have not considered the change in signal strength with time following excitation. In GE imaging, the signal decays with a time constant of 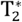 (Figure 1, GE, MTF), which subsumes tissue dependent spin-spin relaxation (time constant *T*_2_) and additional dephasing due to magnetic field inhomogeneities (time constant 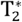). In SE imaging, the signal similarly decays with time constant 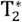. At half the echo time, however, a refocusing pulse causes reversal of the accumulated decay, while *T*_2_ decay continues. After the echo time is reached, the signal returns to a decay with time constant 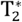 (Figure 1, SE, MTF).

During the decay, the k-space is sampled for the total acquisition time from the smallest (most negative) k-space value to the highest k-space value, such that the center of k-space (k=0) is sampled at the echo time. Note that this common sampling order, also called linear ordering, is not the only one possible. For example, in centric ordering, the sampling trajectory starts at the center of k-space and alternates between increasingly positive and decreasingly negative k-space coordinates. Here, we only consider linear ordering. For each k-space step in the phase-encode direction, an entire line along the read-out direction is acquired while the signal decays only minimally. Consequently, the effect of signal decay on the read-out direction can be neglected. However, in the phase-encode direction, the signal decay modulates the sampled data, causing different weighting of different spatial frequency components. The MTFs of GE and SE along the phase-encoding direction reflect this weighting (Figure 1, GE and SE, MTF). Therefore, the complex PSFs of GE and SE MRI differ from the complex PSF of the imaging process with no signal decay (Figure 1, GE and SE, complex PSF). The time constants used in our simulations reflect imaging at 7 Tesla; unless specifically mentioned otherwise, we considered a total readout duration of 27.8 ms (equal to 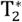 of gray matter). The FWHM of the magnitude PSFs were 1.34 voxels for GE and 1.32 voxels for SE, both larger than the FWHM of the magnitude PSF with no decay (1.20 voxels).

### The Effect of the MRI process on the effective spatial resolution for imaging cortical columns

In the previous section we have discussed why the magnitude PSF does not correctly describe the MR imaging process. Does it follow then, that the FWHM of the magnitude PSF fails to accurately describe the effective spatial resolution of fMRI?

To address this question specifically in the context of imaging cortical columns, we simulated fMRI sampling of BOLD responses of patterns of ocular dominance columns (Chaimow et al., 2011; Rojer and Schwartz, 1990).

Figure 2 (simulated pattern, 2D pattern) shows a 2D modeled pattern and an excerpt from a 1D pattern (simulated pattern, 1D pattern excerpt). The 1D pattern follows the horizontal direction of the 2D pattern, here considered as the phase-encode direction.

**Fig. 2.**
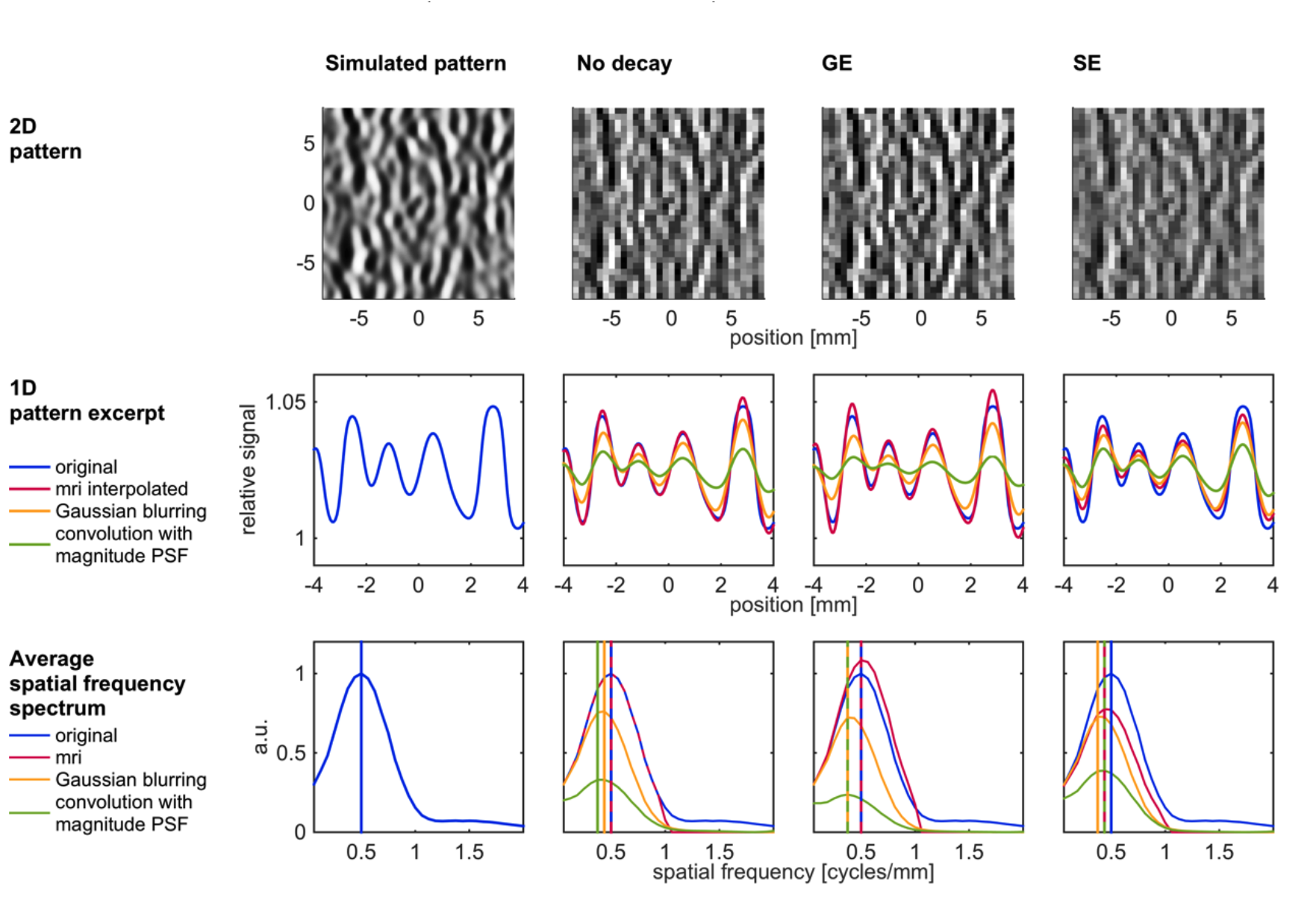
The effect of imaging PSF on MRI of a columnar pattern. We simulated and analyzed columnar ocular dominance patterns (first column) and their MR imaging with no signal decay (second column), GE imaging (third column) and SE imaging (fourth column). The first row shows the simulated original 2D pattern and the resulting MR images using a voxel size of 0.5 mm. The second row shows an extract from a 1D simulated pattern (blue) and its no-decay, GE, and SE imaged counterparts (red). The 1D imaged patterns (which could be presented as non-continuous functions due to voxelization) were interpolated using zero-filling in k-space in order to facilitate comparison. The no-decay imaged pattern was very similar to the original pattern. The GE and SE imaged patterns showed slight increases and decreases, respectively, in contrast. In addition, we simulated convolutions (in orange) of the original pattern with Gaussian PSFs of the same widths as those computed from the magnitude PSFs (black arrows in Fig. 1) and convolutions with the magnitude PSF itself (in green). They all show lower contrast than that in the corresponding MR simulations. The third row shows spatial frequency spectra averaged from 1,000 simulated 1D patterns. The spatial frequency showing the maximal amplitude in each spectrum is marked with a vertical line with corresponding color. In cases where segments of the spectra obtained from the original pattern and the pattern obtained by MRI sampling were identical, we present alternating dashed red (for MRI sampling) and blue (for the original pattern) curves. Similarly, we present alternating dashed blue and red vertical lines in cases for which the frequencies showing the maximal amplitude were identical across the original pattern and MRI sampling. Imaging with no decay did not change the frequency spectrum within the sampled range (up to 1 cycle/mm). For GE, no change in the frequency showing the maximal amplitude was detected at the resolution we applied. SE imaging resulted in a slightly lower spatial frequency showing the maximal amplitude. In contrast, all spectra obtained by the Gaussian convolution (orange) and by the magnitude PSF convolution (green) showed lower spatial frequencies associated with the maximal amplitudes.

These 2D and 1D modeled patterns represent a BOLD pattern, consisting of a spatially constant baseline signal of 1 and a superimposed ODC pattern-dependent BOLD response. The BOLD response can vary between 0% and 5% relative to the baseline value. We did not model the spatial spread of the BOLD response (meaning we assumed no spread) in order to not distract from the effects of the imaging process.

The main spatial frequency of the simulated ODC BOLD pattern was 0.5 cycles/mm (Yacoub et al., 2007), reflected as the maximum in the spatial frequency spectrum (Figure 2, simulated pattern, average spatial frequency spectrum, vertical blue lines). The irregularity of the pattern is reflected in a distribution of additional spatial frequency contributions around the two maxima.

In order to quantify the functional contrast of true or imaged responses, we defined the contrast range as the standard deviation around the average response (the average response is defined as one half of the maximum response relative to the background intensity, namely 1.025). The contrast range averaged over 1,000 simulated one-dimensional patterns, was 1.30% (relative to the baseline of 1).

### MRI with no signal decay

First we analyzed the effect of MR imaging with no signal decay. We simulated MRI sampling of the simulated original pattern (Figure 2, Simulated pattern) using a voxel width of 0.5 mm. Figure 2 (No decay, 2D pattern) shows the corresponding imaged two-dimensional pattern. Figure 2 (No decay, 1D pattern excerpt) shows the values of the imaged and interpolated one-dimensional pattern (red). The MR imaged pattern is shown as a high-resolution representation, which is equivalent to an interpolation using zero-filling in k-space. This was done in order to facilitate visual comparison to the original pattern (the alternative presentation format would consist of a discrete function due to voxelization). The imaged pattern was very similar to the original pattern (blue). The average contrast range computed over 1,000 imaged patterns decreased only slightly from 1.30% (average SD of 1,000 original patterns) to 1.29% (average SD of 1,000 no-decay MRI patterns).

In addition, we compared the average frequency spectrum of the imaged patterns to the average spectrum computed over the original patterns (Figure 2, no decay, average spatial frequency spectrum). Within the sampled k-space range (spatial frequencies below 1 cycle/mm), the two spectra were identical. Outside of this range, the average spectrum of the imaged patterns was zero.

Note, however, that the FWHM of the magnitude PSF corresponding to MRI with no signal decay was 1.20 voxels (= 0.6 mm in our specific simulation). This result could be wrongly interpreted to imply that MR image formation, ignoring decay, is comparable to blurring with a kernel (e.g. a Gaussian) of the same width.

Figure 2 (no decay, 1D pattern excerpt, orange curve) shows that in contrast to actual MRI sampling, such blurring would have resulted in a reduced amplitude (contrast range = 0.95%) and a shift in the spatial frequency spectrum to 0.4375 cycles/mm (Figure 2, no decay, average spatial frequency spectrum, orange lines). Using the magnitude PSF as a convolution kernel in itself (although, in fact, it is not a convolution kernel of the MRI process) resulted in an even larger reduction in contrast (0.45%, Figure 2, no decay, 1D pattern excerpt, green) and a shift of the spatial frequency distribution towards lower frequencies (maximum at 0.3750 cycles/mm, Figure 2, no decay, average spatial frequency spectrum, green).

### MR imaging in the presence of signal decay: Gradient-Echo imaging

Next, we analyzed the effect of signal decay. Figure 2 (GE, 2D pattern) shows the GE imaged two-dimensional pattern. There is no noticeable blur relative to the no-decay image. In fact, the interpolated one-dimensional imaged pattern (Figure 2, GE, 1D pattern excerpt, red) showed a higher amplitude compared to the original pattern (blue). The average contrast range increased from 1.30 % (original) to 1.41% (GE).

This increase in contrast did not result in a noticeable difference in the peak spatial frequency, which remained at 0.5 cycles/mm (Figure 2, GE, average spatial frequency spectrum; the resolution we employed in our simulated k-space was 0.0625 cycles/mm). However, relative to the average spectrum of the original patterns, the amplitude at the peak spatial frequency increased, while the amplitudes at spatial frequencies close to 0 cycles/mm remained constant. This shows that GE imaging had the effect of a moderate high-pass filter. Similar to our conclusion for the imaging with no signal decay, this result could not be expected by simply considering the positive FWHM of the magnitude PSF, which was 1.34 voxels (= 0.67 mm in our specific simulation).

A convolution with a Gaussian of the same width resulted in contrast reduction (Figure 2, GE, 1D pattern excerpt, orange). The average contrast range dropped from 1.30% to 0.89%. Also, the peak in the spatial frequency spectrum shifted to a lower frequency of 0.375 cycles/mm. Convolution of the original pattern with the magnitude PSF resulted in a larger reduction in contrast (average contrast range 0.33%) and a shift of the peak spatial frequency towards lower frequencies (0.375 cycles/mm).

### MR imaging in the presence of signal decay: Spin-Echo imaging

Figure 2 (SE, 2D pattern) shows the SE imaged two-dimensional pattern. The SE image is slightly blurred relative to the image obtained with no signal decay. Similarly, the interpolated one-dimensional pattern excerpt (Figure 2, SE, 1D pattern excerpt, red) shows a lower amplitude compared to the original pattern (blue). The average contrast range decreased from 1.30% (original) to 1.01% (SE) and the peak spatial frequency shifted to a lower frequency of 0.4375 cycles/mm (Figure 2, SE, average spatial frequency spectrum).

The reduction in contrast and peak spatial frequency suggests that SE imaging had a blurring effect on the original pattern, which could be consistent with the spatial extent of its magnitude PSF (FWHM of 0.66 mm). However, convolution with a Gaussian of the same width resulted in even larger contrast reductions (Figure 2, SE, 1D pattern excerpt, orange). The average contrast range decreased further to 0.90% and the peak in the spatial frequency spectrum shifted to 0.375 cycles/mm. True also for SE, convolution with the magnitude PSF reduced the contrast (0.52%) more than the MRI simulation and the convolution with a Gaussian kernel (of the same width as the width of the magnitude PSF) did. The peak spatial frequency remained at 0.4375 relative to the SE MR simulation.

### An alternative approach to quantifying the effect of MR imaging on the effective spatial resolution

#### Convolution with the real component of the complex imaging PSF linearly approximates the complete MRI process

We have shown that neither the linear process of convolution with the complex PSF (without taking the absolute) nor the linear process of convolution with the magnitude PSF can faithfully describe the entire non-linear MR imaging and reconstruction process. To characterize the complete MRI process using a PSF, we propose an alternative, optimal, linear approximation.

The best linear approximation of a function around a point *x*_0_ is the derivative of the function at *x*_0_. For a function of a single variable *f*(*x*), the derivative *f*′(*x*) represents a tangent line which can be interpreted as a linear approximation to *f*(*x*) by mapping small deviations *x*_0_ + Δ*x* onto 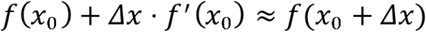.

In the case of the MR imaging and reconstruction process, the function under consideration is not a function of a single variable but a functional, which maps a pattern onto a set of imaged voxel values. The derivative of this functional is a linear transformation that itself depends on a baseline pattern (the *point* at which the derivative is evaluated) in the same manner that a tangent depends on the point (*x*_0_) at which it is defined. Similar to a tangent line, this linear transformation maps a pattern with small deviations from the baseline pattern onto a set of voxel values approximating the true imaged pattern.

In Appendix B, we computed this derivative for a spatially constant baseline pattern. This derivative is a linear transformation that approximates the complete MR process for other patterns, with small response deviations relative to this spatially constant baseline pattern. In appendix B we show that this linear approximation is identical to a convolution with the real component of the complex imaging PSF (presented in the middle row of Figure 1, in purple)

Next, we evaluated how well this convolution approximates the true MR imaging and reconstruction process. In particular, because the linear approximation is expected to be valid for small deviations around the spatially constant baseline pattern, we quantified the dependence of the quality of the approximation on the response amplitude. To this end, we simulated a wide range of columnar patterns with different response amplitudes. We then compared the results obtained by simulating the full MRI process for each of these patterns to those obtained by a convolution of the pattern with the real component of the complex imaging PSF (Fig. 3). Note that as expected, for large amplitude deviations from the spatially constant baseline pattern, the linear approximation can result in relatively large errors (e.g. 95th percentile of the RMSE were 6.13% and 0.68% for 100% response amplitude imaged with GE and SE fMRI, respectively; Figure 3, upper row). However, the root-mean-squared error for realistic response amplitudes was small. For example, for the maximal response amplitude of 5% imaged with GE fMRI, the 95th percentile was 0.43% of the standard deviation of the pattern obtained by the full MRI simulation (Figure 3, middle row). For SE imaging, the relative RMSE was even lower; for the maximal response amplitude of 5% imaged with SE fMRI it was 0.05% f (Figure 3, middle row). In contrast, convolution of the same response patterns (with response amplitude of 5%) with the magnitude PSF resulted in median relative root-mean-squared errors of 78% (GE) and 48% (SE) (Figure 3, bottom row).

**Fig. 3.**
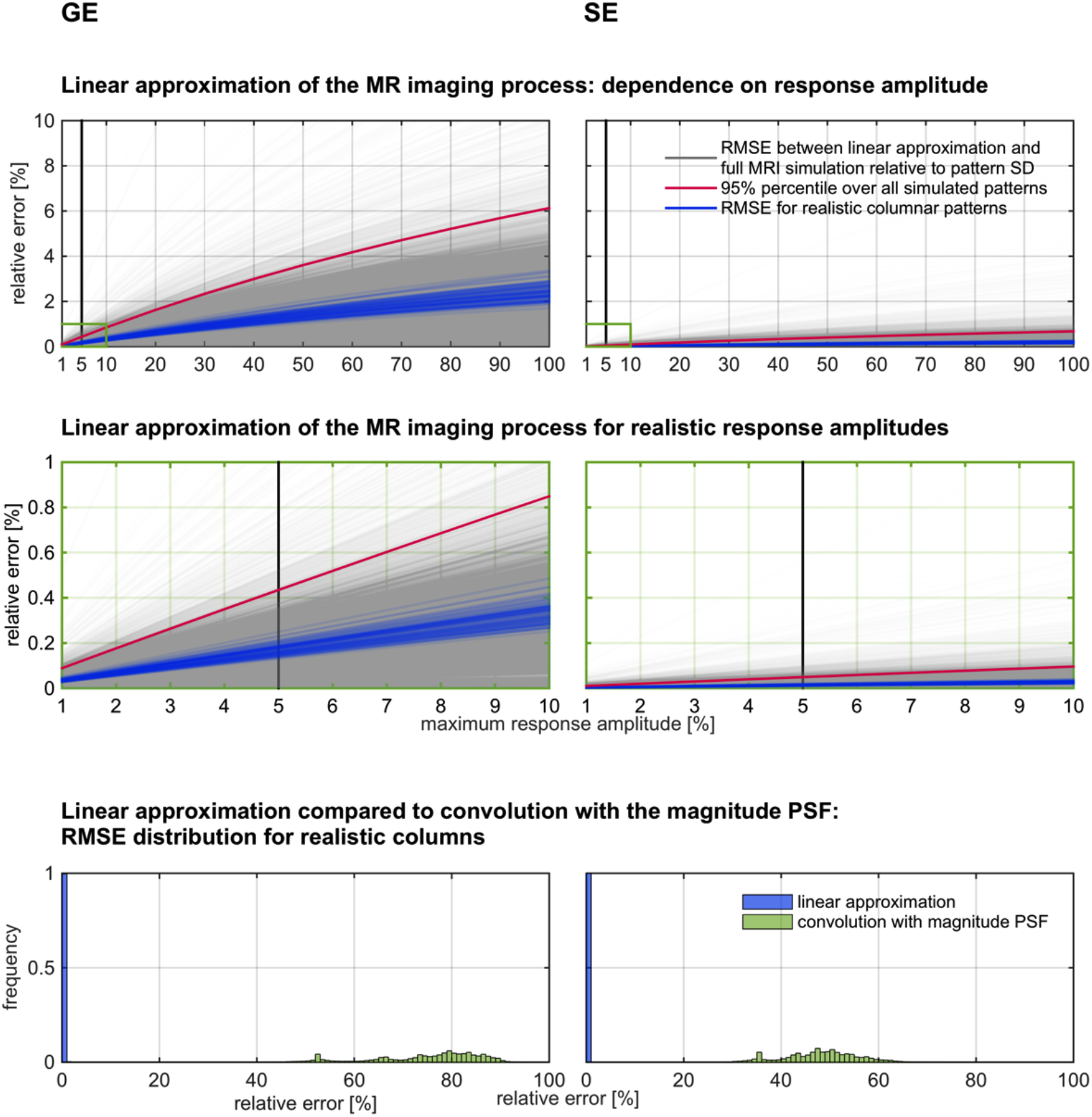
Evaluation of the linear approximation of the complete MRI process. The upper row shows the dependence of the linear approximation for GE imaging (left) and SE imaging (right) as a function of maximum response amplitude. Cortical columnar response patterns for a wide range of spatial frequency parameters, irregularity parameters, and maximum response amplitudes were simulated (1,000 for each parameter combination; results from 100 patterns were used for visualization). MR imaging of these patterns was simulated. In addition, a linear approximation of the MR imaging process consisting of a convolution of the pattern with the real component of the complex imaging PSF was computed. The results of the linear approximations were compared to the complete MR imaging simulations by means of the root-mean-squared-errors (RMSE) relative to the standard deviation of the images obtained by the complete MRI process. The gray curves show the distribution of relative RMSEs from all simulated patterns; the red curve presents their 95 percentile. The blue curves show results from realistic columnar parameters (intermediate irregularity and spatial frequency). All curves show increased errors with increasing response amplitude. For the default, realistic response amplitude level of 5% (indicated by a vertical black line), most relative errors were well below 1%. The middle row of panels presents a magnified view of the errors for such realistic response amplitudes. The bottom part of the figure compares the distribution of relative RMSEs obtained by the linear approximation with those obtained by a convolution with the magnitude PSF. The default, realistic response amplitude level of 5% was used for this comparison. The convolution with the magnitude PSF resulted in substantially higher errors.

### Quantifying the effect of signal decay by fitting a two-component model consisting of convolution with a Gaussian PSF followed by MR sampling with no decay

In itself, the real component of the complex imaging PSF, in particular its width, is not suited to characterize the effective spatial resolution of the MR imaging process. The reason is that it represents not only signal decay, but also the MR sampling process. As we have shown, the latter is pattern-dependent and irrelevant if the voxels are sufficiently small to sample the spatial frequency spectrum, such that the imaged pattern is similar to the original pattern. Furthermore, it is not possible to easily discriminate the blurring characteristics of SE from the high-pass filtering characteristics of GE on the basis of the real component of the complex imaging PSFs.

However, instead of considering the real components of the complex imaging PSFs, we can consider their spatial frequency representations. Figure 4 (second row) shows the MTF of the real component of the complex imaging PSF for GE and SE (blue). The MTF of the real component is equal to the average of the positive and negative components of the original imaging MTF (presented in Figure 4, first row), assigned in a mirror-symmetric manner around the center (k=0) of the k-space.

**Fig. 4.**
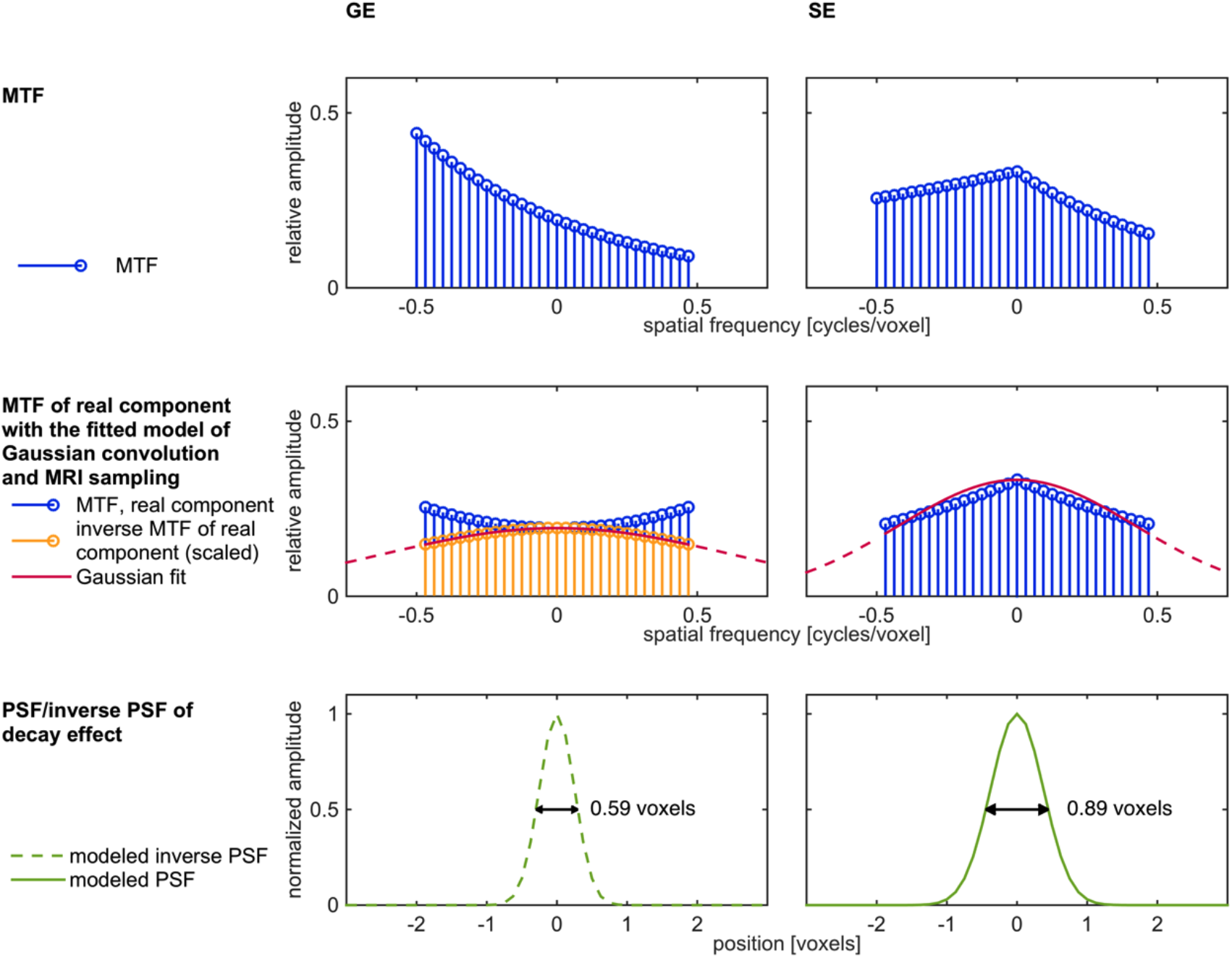
Fitting of a two-component model consisting of convolution with a Gaussian PSF that accounts for signal decay followed by MR sampling with no decay. This figure shows the fitting of a Gaussian convolution and MRI sampling model for GE imaging (first column) and SE imaging (second column). Gaussian functions (second row, red) were fitted to the MTF of the real component of the SE image PSF (second row, right side column, in blue) and to the scaled inverse MTF of the real component of the GE image (second row, left side column, in orange). Outside the sampled k-space range, the continuation of the Gaussian fit is shown as a dashed red line. The Fourier transforms of these functions are Gaussian PSFs (bottom row, in green). For SE, this PSF describes the blurring due to the signal decay. For GE, it describes a hypothetical blurring that would be reversed by the high-pass filter properties of the 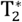 decay effect. Their respective FWHM (black arrows) are 0.59 voxels (GE) and 0.89 voxels (SE).

The MTF of the real component of the complex imaging PSF can be regarded as the product of two factors representing two different processes. The first factor is the rect-function-windowed Dirac comb function that describes the MRI sampling with no signal decay. The second factor is a modulation of the sampled signal due to signal decay. In order to model this second factor we can choose a function that fits the modulation within the sampled k-space range. The Fourier transform of this function will be a convolution kernel in the image space, which describes the effect of signal decay. It does not describe the effect of MRI sampling with no signal decay, which is qualitatively different and depends on the imaged pattern.

We can then apply MRI sampling with no signal decay. The image we get from the convolution that accounts for the signal decay and the MRI sampling is either identical to the complete MRI process (depending on the function fitted to the k-space representation in its sampled range) or approximates it.

For SE imaging, we chose a Gaussian function and fitted it to the signal decay-dependent modulation of the MTF of the real component of the complex imaging PSF (= the second factor; Figure 4 second row, SE, red; See discussion for justification of modeling the second factor as a Gaussian function). An inverse Fourier transformation of this fitted Gaussian results in a Gaussian PSF in the image space, allowing for the interpretation of the signal decay effect as Gaussian blurring. The FWHM of this Gaussian PSF was 0.89 voxels (for a total read duration of 27.8 ms at 7T).

For GE imaging, a Gaussian function is not a good fit, since the MTF of the real component of the complex imaging PSF shows increasing amplitudes with increasing spatial frequency (Figure 4, second row, GE, blue), consistent with its high-pass filtering properties we have shown above (Figure 2). However, instead of this MTF, we can consider its inverse (1 divided by the MTF; Figure 4, second row, GE, orange). The inverse MTF describes the process that would be reversed by a convolution with the real component of the complex PSF. We fitted a Gaussian to the signal decay-dependent modulation of this inverse MTF (which resulted in a higher R^2^, than the Gaussian fit to the non-inverted MTF) and calculated its corresponding Gaussian PSF in the image space. This allows for interpreting GE imaging as reversing (deconvolving) a Gaussian blur. The FWHM of this Gaussian was 0.59 voxels (for a total read duration of 27.8 ms at 7T).

### Partial Fourier acquisition

In addition to the standard EPI acquisition described so far, one can shorten the total read-out duration by only acquiring parts of the conjugate symmetric k-space. This is known as partial Fourier acquisition.

In order to study how signal decay affects partial Fourier acquisition, we simulated MTFs resulting from partial Fourier imaging in which either the first ¼ or last ¼ of the phase-encode steps were omitted (the total read out duration was shortened accordingly from 27.8 ms to 20.85 ms). We then applied the same modeling methodology described above for the full k-space acquisitions (Figures 5 and 6 for GE and SE, respectively). In addition, we compared a reconstruction that exploited the conjugate symmetry (Figures 5 and 6, columns 1 and 3) to a reconstruction with simple zero-filling (Figures 5 and 6, columns 2 and 4).

For GE, partial Fourier with early omission resulted in blurring (Figure 5 columns 1 and 2) as opposed to the high-pass filtering observed in full k-space acquisition (Figure 4). The blurring was more substantial in zero-filling reconstruction (FWHM = 1.38 voxels) than in conjugate symmetry reconstruction (FWHM = 1.00 voxels).

**Fig. 5.**
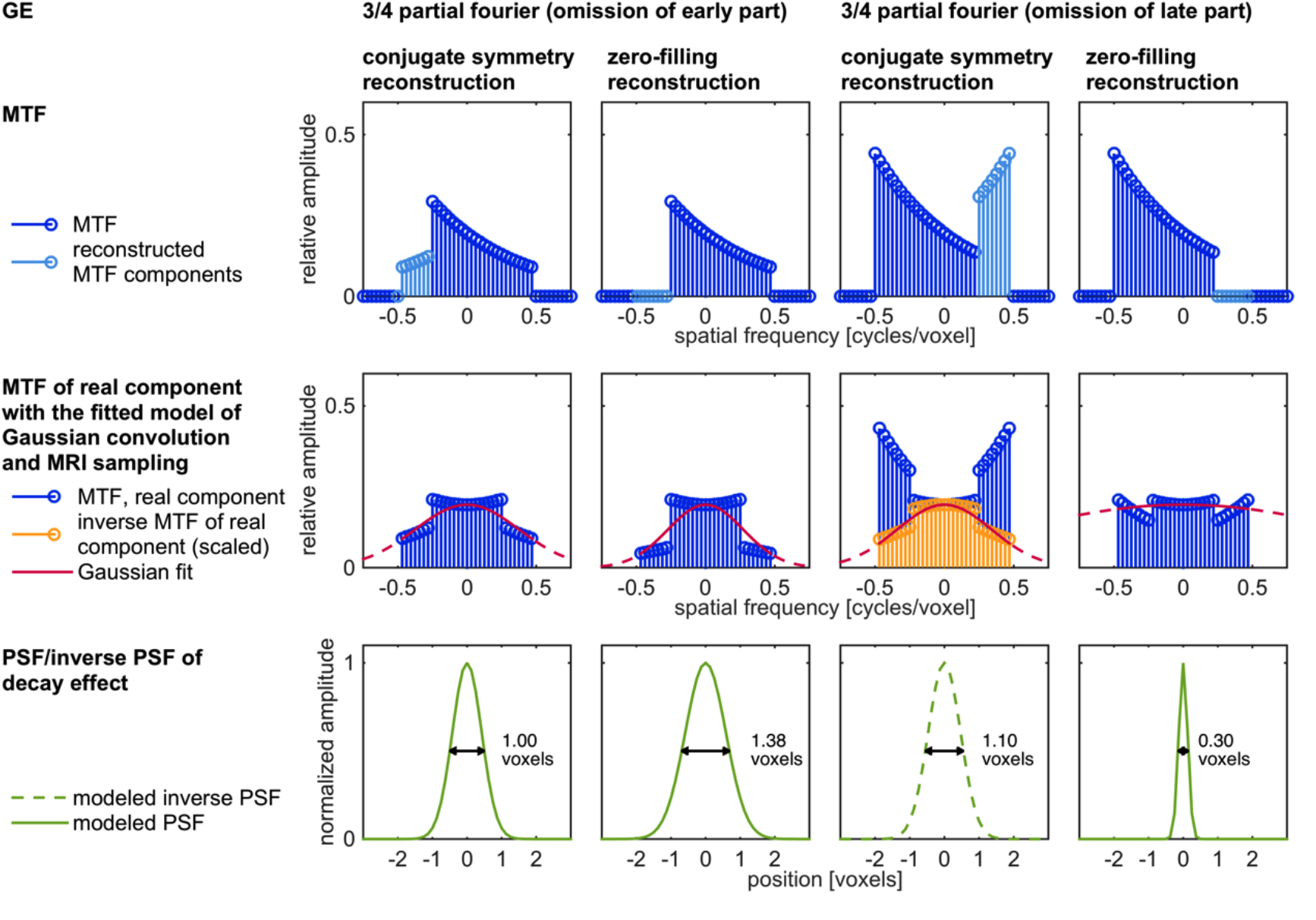
Fitting of the two-component model for GE partial Fourier acquisition. This figure shows the fitting of a two-component model for partial Fourier acquisition using GE imaging. Omission of the first ¼ (columns 1 and 2) and last ¼ (columns 3 and 4) of phase-encode steps were simulated. Furthermore, a reconstruction that exploits conjugate symmetry (columns 1 and 3) was compared to a zero-filling reconstruction (columns 2 and 4). The first row shows the imaging MTF resulting from measurement components (dark blue) and reconstruction components (light blue). Gaussian functions (second row, in red) were fitted to the MTF of the real component of the imaging PSF (early-omission partial Fourier, columns 1 and 2, and late-omission partial Fourier using zero-filling reconstruction, column 4, in blue) and to the inverse MTF (here scaled for clarity of presentation) of the real component of the imaging PSF (late-omission partial Fourier using conjugate symmetry reconstruction, column 3, in orange). Outside the sampled k-space range, the continuation of the Gaussian fit is presented as a dashed red line. The Fourier transforms of these functions are Gaussian PSFs (bottom row, in green). For early-omission partial Fourier (columns 1 and 2) and late-omission partial Fourier using zero-filling reconstruction (column 4), these PSFs describe the blurring due to the signal decay. For late-omission partial Fourier using conjugate symmetry reconstruction (column 3), the PSF describes a hypothetical blurring that would be reversed by the high-pass filter properties of the 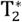 decay effect.

Partial Fourier with late omission using conjugate symmetry reconstruction resulted in high-pass filtering (Figure 5, column 3). This high-pass filtering effect (reverse kernel FWHM = 1.10 voxels) increased relative to that of full k-space acquisition (reverse kernel FWHM = 0.59 voxels). Partial Fourier with late omission using zero filling reconstruction (Figure 5, column 4) resulted in moderate low-pass filtering (FWHM = 0.30 voxels).

For SE, partial Fourier with early omission (Figure 6, columns 1 and 2) resulted in increased blurring relative to that shown by the full k-space acquisition (FWHM = 0.89 voxels). The blurring was more substantial in zero-filling reconstruction (FWHM = 1.55 voxels) than in conjugate symmetry reconstruction (FWHM = 1.10 voxels). Partial Fourier with late omission also resulted in blurring (Figure 6, columns 3 and 4). Here, conjugate symmetry and zerofilling reconstructions resulted in decreased (FWHM = 0.66 voxels) and increased (FWHM = 1.38 voxels) blurring, respectively, relative to the blurring obtained from the full k-space acquisition (FWHM = 0.89 voxels).

**Fig. 6.**
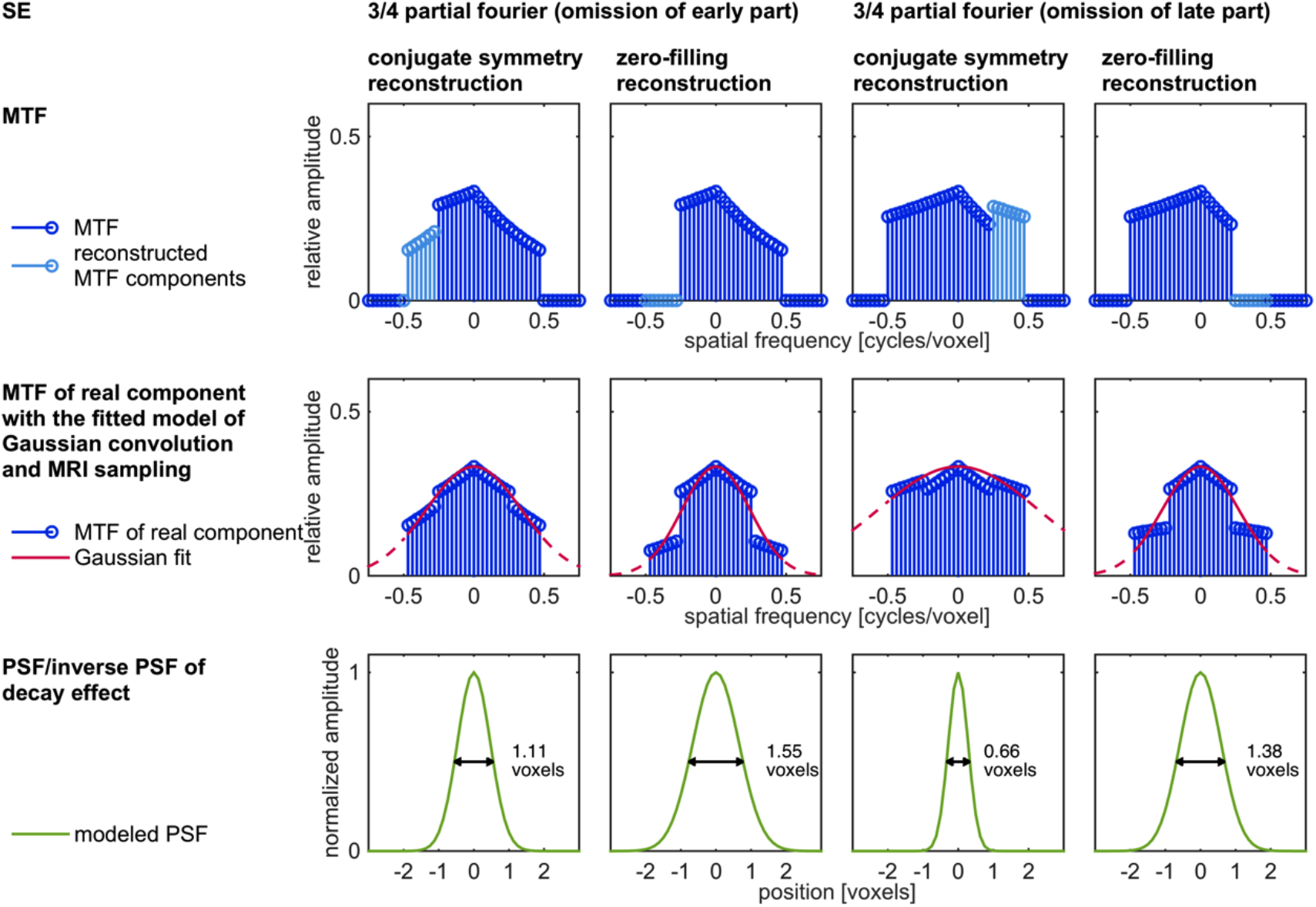
Fitting of the two-component model for SE partial Fourier acquisition. This figure shows the fitting of a two-component model for partial Fourier acquisition using SE imaging. Omission of the first ¼ (columns 1 and 2) and last ¼ (columns 3 and 4) of phase encode steps were considered. Furthermore, a reconstruction that exploited conjugate symmetry (columns 1 and 3) was compared to a zero-filling reconstruction (columns 2 and 4). The first row shows the imaging MTF resulting from measurement components (dark blue) and reconstruction components (light blue). Gaussian functions (second row, in red) were fitted to the MTF of the real component of the imaging PSF for all columns 1-4. Outside the sampled k-space range, the continuation of the Gaussian fit is presented as a dashed red line. The Fourier transforms of these functions are Gaussian PSFs (bottom row, in green) that have a blurring effect. These PSFs describe the blurring due to the signal decay.

All results so far were based on an assumed total readout duration of 27.8 ms. In order to extend our results and to study the dependence of signal decay blurring on the total read out duration, we repeated the simulation of signal decay-dependent MTFs and the fitting of our model of Gaussian convolution and MRI sampling for a range of total readout durations (Figure 7).

**Fig. 7.**
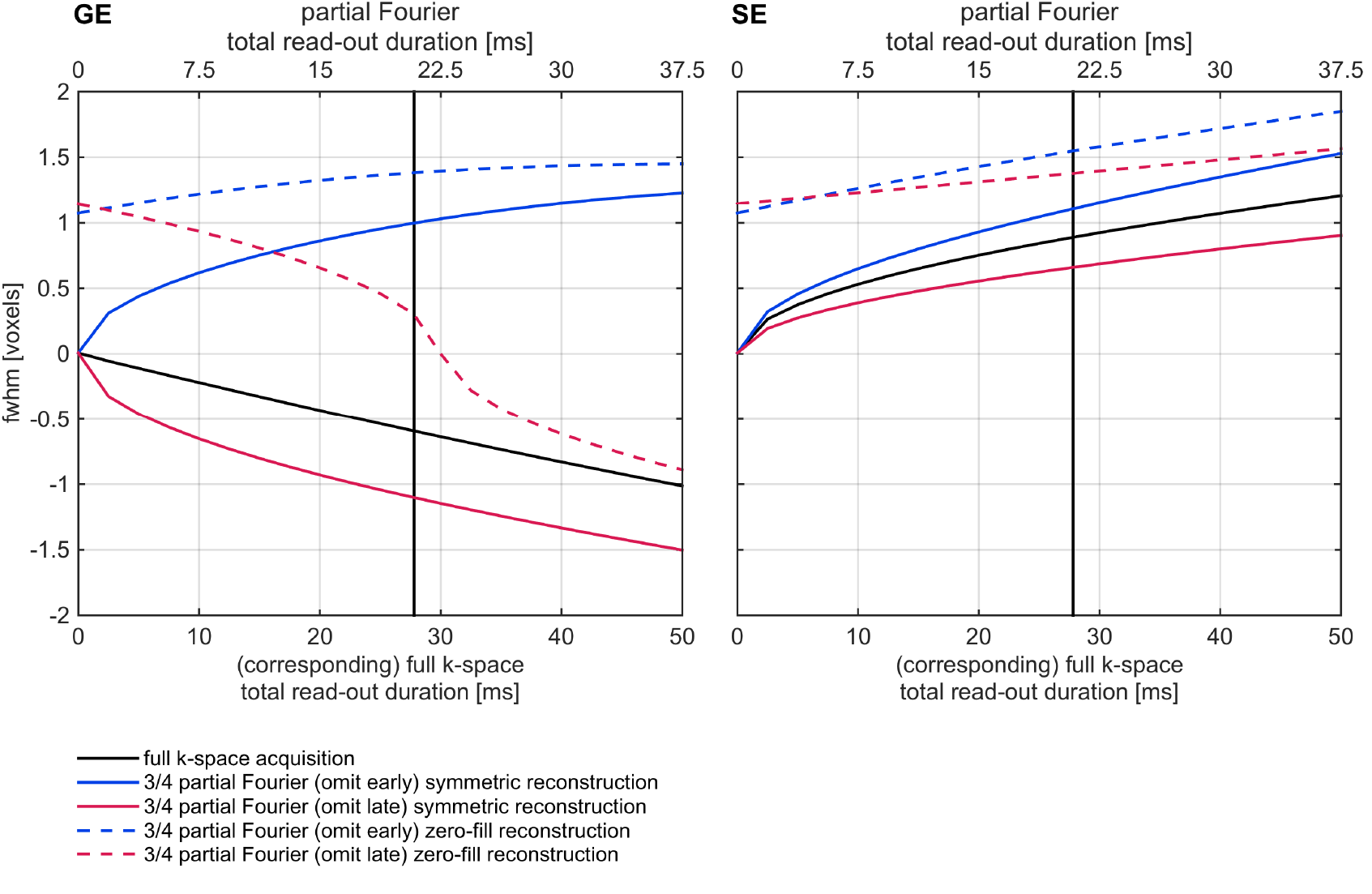
FWHMs of Gaussian PSFs that model the effect of signal-decay as a function of total read-out duration. This figure shows the results of fitting our two component model that accounts for MR sampling and signal decay for different GE (left) and SE (right) imaging scenarios and for different total read-out durations. The FWHM of the fitted Gaussian PSFs are presented as a function of total-read out duration. For Partial Fourier acquisition, the total-read out duration axis is labeled at the top. Partial Fourier acquisition total-read out durations were shortened according to the fraction of omitted k-space (1/4) relative to the corresponding full k-space total read-out duration (bottom axis). The vertical black line represents a total-read out duration of 27.8 ms (20.85 ms for partial Fourier). Negative FWHM values indicate that the Gaussian PSFs resulted from the inverse of the MTF of the real component of the complex imaging PSF. Such negative values represent a hypothetical blurring that is reversed by the high-pass filter properties of the 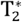 decay effect.

For almost all imaging scenarios, including GE and SE imaging, the type of decay-dependent effect (blurring or high-pass filtering) was independent of total readout duration. However, the effect s strength increased with increasing total readout duration. The only exception was the zero-filling reconstruction of GE partial Fourier imaging that omits the late acquisitions. Here, short total readout durations resulted in blurring while longer total readout durations resulted in high-pass filtering.

### Evaluation of modeling the complete MR process as a convolution with a Gaussian PSF that accounts for signal decay followed by MR sampling with no decay

Lastly, we evaluated how well our simplifying model approximated a complete MRI acquisition model (Figure 8). We compared complete simulations of fMRI of columnar patterns including signal decay to simulations that used our Gaussian PSF model of signal decay followed by MR sampling with no decay. We then quantified their deviations by calculating the root-mean-squared errors relative to the standard deviation of the patterns obtained by the simulation of the complete MRI process. The relative errors differed for different imaging methods and total readout durations. For full k-space acquisition (Figure 8, upper row), the median error (red curve) obtained for a total readout duration of 27.8 ms was 3.91% and 8.39% using GE and SE fMRI, respectively.

**Fig. 8.**
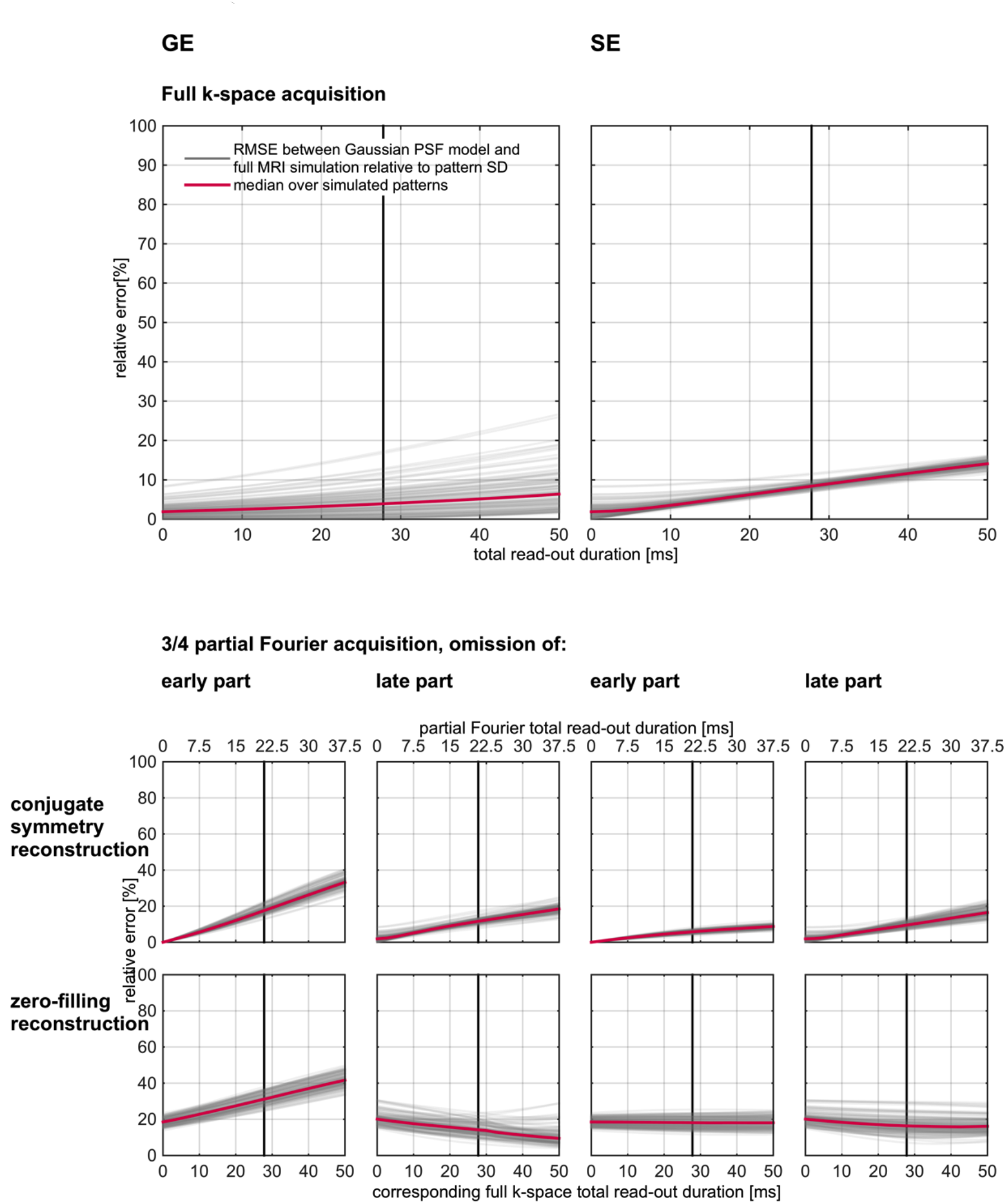
Evaluation of modeling the complete MR process as a convolution with a Gaussian PSF that accounts for signal decay followed by MR sampling with no decay. 1,000 cortical columnar response patterns (with intermediate irregularity and spatial frequency) were simulated (results from 100 patterns were used for visualization). For GE (left) and SE (right) full k-space acquisition (top) and partial Fourier acquisition (bottom), convolution with the fitted Gaussian PSF (or its inverse model, see text) followed by MR sampling with no decay was compared to the complete MR imaging process. To this end, the root-mean-squared-errors (RMSE) relative to the standard deviation of the complete MR images were calculated. The gray curves show the distribution of relative RMSEs from all simulated patterns as a function of total read-out duration. The vertical black line represents a total-read out duration of 27.8 ms (20.85 ms for partial Fourier). The medians of RMSEs are shown in red. For Partial Fourier acquisition, the total-read out duration axis is labeled at the top. Partial Fourier acquisition total-read out durations were shortened according to the fraction of omitted k-space (1/4) relative to the corresponding full k-space total read-out duration (bottom axis). RMSEs were generally low for full k-space acquisition but became higher for the majority of partial Fourier acquisition schemes.

The relative errors obtained for the majority of partial Fourier acquisition schemes were substantially higher. For conjugate symmetry reconstruction (Figure 8, middle row), the median error computed for GE fMRI and a total readout duration of 20.85 ms (that corresponds to 27.8 ms for full k-space acquisition) was 17.55% and 11.6% for omissions of the early and late part of the k-space, respectively. The corresponding errors for SE fMRI were 5.79% and 9.51%.

For zero filling reconstruction (Figure 8, bottom row), the median error computed for GE fMRI and a total readout duration of 20.85 ms was 31.15% and 14.19% for omissions of the early and late part of the k-space, respectively. The corresponding errors for SE fMRI were 18.14% and 16.37%.

Our Gaussian PSF model of signal decay approximates the signal decay effect in MR imaging as a pattern-independent linear process. As we have just demonstrated, the differences between the results of this approximation and the true imaging process are low for full k-space acquisitions, but are higher for partial Fourier acquisitions. In order to obtain an even better approximation, we can define an alternative approximation. Given a specific pattern, we can determine a Gaussian PSF (or its inverse process) that accounts for signal decay, such that convolution with this specific Gaussian followed by MR sampling with no decay results in the best Gaussian-based approximation of the complete MR imaging process.

Figure 9 presents the resulting Gaussian PSF FWHMs (blue, median in red) as a function of total read-out duration for full k-space acquisitions (upper row) and for different partial Fourier acquisition schemes (middle and bottom rows). In addition, we present the previously estimated pattern-independent Gaussian PSF FWHMs (dashed orange curves). Note that the patterns used for this evaluation were the results of different columnar patterns, but they all shared the same statistical properties (main pattern frequency = 1 cycle per 4 voxels, relative irregularity = 0.5).

**Fig. 9.**
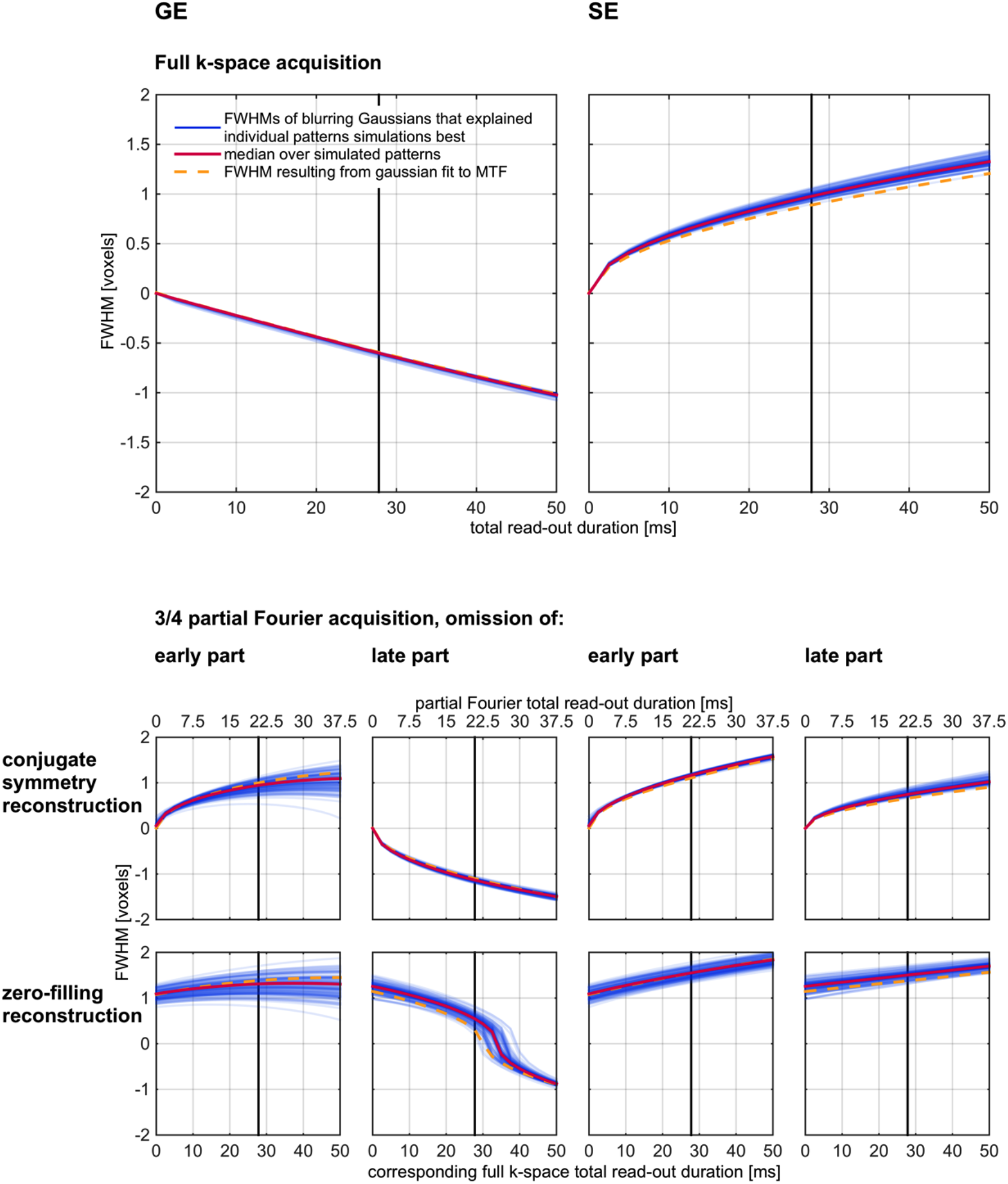
Estimation of pattern specific Gaussian PSFs that model the effect of signal-decay. 1,000 cortical columnar response patterns (with intermediate irregularity and spatial frequency) were simulated (results from 100 patterns were used for visualization). We present results from GE (left) and SE (right) full k-space acquisition (top) and partial Fourier acquisition (bottom). For each pattern, complete MR imaging was simulated and compared to the result of convolution of the pattern with Gaussians (or their inverse models, see text) followed by MR sampling with no decay. For each pattern, the pattern specific Gaussian that resulted in the smallest mean squared error relative to the complete MR imaging simulation were determined. The blue curves show the distribution of FWHMs of these Gaussians as a function of total read-out duration. The vertical black line represents a total-read out duration of 27.8 ms (20.85 ms for partial Fourier). The medians of FWHMs are shown in red. For Partial Fourier acquisition, the total read-out duration axis is labeled at the top. Partial Fourier acquisition total read-out durations were shortened according to the fraction of omitted k-space (1/4) relative to the corresponding full k-space total read-out duration (bottom axis). The FWHMs of our pattern independent Gaussian PSF model are presented in dashed orange for comparison.

The pattern-dependent PSF widths (blue curves) for the full k-space acquisitions did not vary much across patterns (they were relatively independent of the specific pattern). They corresponded reasonably well to those obtained from our pattern-independent model (dashed orange curve). For the majority of partial Fourier acquisition schemes, the variability of pattern-specific PSF widths was somewhat higher. In addition, the differences between the median over simulated patterns and the pattern independent model were on average higher than those obtained for the full k-space acquisitions. This suggests that using a pattern-independent, single PSF width does not fully characterize the effective spatial resolution under partial Fourier acquisitions. We note, however, that for both full k-space and partial Fourier acquisitions, the estimations obtained from our proposed pattern-independent model (in dashed orange curves) matched those obtained from the pattern-dependent simulations reasonably well.

## Discussion

### The Imaging PSF and effective spatial resolution

The PSF of an imaging system is defined as the image obtained from an infinitesimally small point-like object. If the imaging system is linear and shift invariant, its response to an arbitrary object can be described as a convolution with the imaging PSF as the convolution kernel. This latter property is what makes the PSF useful in describing the spatial characteristics of an imaging system.

The effect of a system with an imaging PSF with its maximum at zero and whose strictly positive values do not increase with increasing distance from zero can be intuitively understood as spatial spreading or blurring. Such systems produce smoothed image versions of any object or pattern. The smoothing can be quantified by measures of the imaging PSF width (such as the FWHM). However, other PSF shapes may characterize certain systems, resulting in more complex effects that require careful evaluation and may not be intuitive.

Note that the width of the PSF is not the only possible measure of effective spatial resolution. Another measure that has been proposed is the area under the PSF divided by the value of the PSF at the origin (Haacke et al., 1999). In the case of MR imaging without signal decay, it is identical to the actual voxel width.

### The magnitude PSF is not a good measure for the effective spatial resolution of MRI and fMRI

We have shown that the FWHM of the magnitude PSF is not a good measure for the effect of *T*_2_ and 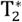 signal decay on spatial resolution. There are three reasons as to why this is the case.

### The larger part of the FWHM of the magnitude PSF is due to MR sampling and not due to signal decay

The magnitude PSF of MR imaging, even with no signal decay, has a FWHM of 1.2 voxels compared to 1.4 voxels and 1.3 voxels (examples given for total readout of 30 ms at 7T) for GE and SE decay, respectively. However this MR sampling effect with no signal decay cannot be regarded as a simple spread of signal. In fact, it acts as a hard low-pass filter that discards all spatial frequency components higher than the voxel size-dependent highest sampled spatial frequency, and leaves all other spatial frequency components unchanged. For a pattern dominated by the latter spatial frequency components, MR sampling with no signal decay has virtually no blurring effect. For a pattern with spatial frequencies limited to those sampled by the MRI process, MR sampling with no signal decay has no blurring effect at all. Thus, the FWHM of the magnitude PSF is a pattern-independent measure, whereas the effective resolution of MRI sampling with no signal decay does depend on the imaged pattern.

### The MR imaging process is non-linear

The last step of the reconstruction in an MR imaging process typically involves taking the absolute values (magnitude image) of the complex image values. The complex image values are the results of a process that can be described as a convolution of the original pattern with the complex imaging PSF. Taking the absolute values of the complex image values makes the MRI process nonlinear. In general, the result of convolving a pattern with a complex kernel, then taking the magnitude image, is not equal to convolving the same pattern with the magnitude values of the complex kernel. This general statement applies in the specific case of MRI: the image obtained by the MRI process is different from that obtained by convolving the original pattern with the magnitude PSF.

To illustrate the effect of the non-linearity of the MRI process, we will describe two scenarios involving the imaging of a point-like structure. While the FWHM of the magnitude PSF can describe the effective spatial resolution of imaging an infinitesimally small point-like structure with no background, it cannot describe the MRI sampling of any arbitrary structure. For example, MRI of a pattern composed of a similar infinitesimally small point-like structure superimposed on a spatially constant baseline with an amplitude higher than the amplitude of the magnitude PSF would include negative side lobes relative to the baseline. Thus, the magnitude PSF fails to correctly describe the MRI process.

Note that even convolving the original pattern with the complex imaging PSF does not fully describe the MRI process, since it does not reflect the operation of taking the magnitude image. Therefore, even the complex imaging PSF on its own does not completely reflect the typical MRI sampling and reconstruction process.

### Signal decay may cause blurring or high-pass filtering with identical FWHM of the magnitude PSF

Signal decay does not always blur the pattern; it can also cause high-pass filtering. This is the case for the GE simulations we have conducted. Whether signal decay results in blurring or high-pass filtering depends on the shape of the decay curve and on the ordering of k-space acquisitions. Specific decay curves and ordering of k-space acquisitions may result in imaging PSFs that are different, but share the same FWHM. Indeed, the magnitude PSFs that are associated with imaging processes that result in blurring or high-pass filtering can be different but may have identical widths. This clearly limits the interpretation based on the magnitude PSF s width measure.

### What does the magnitude PSF describe?

In the 3 previous sub-sections, we have shown that the magnitude PSF is not a good measure for characterizing the effective spatial resolution of the MRI process. The magnitude PSF carries less information relative to the complex imaging PSF. Simply relying on the FWHM of the magnitude PSF further reduces the available information. The magnitude PSF cannot differentiate between reduced effective spatial resolution due to the MRI sampling per se or due to signal decay. The effect of MRI sampling with no signal decay depends on the original pattern, whereas the FWHM of the magnitude PSF does not. While the magnitude PSF does describe the magnitude MR image of a small point-like structure with no background signal, it is not a convolution kernel of the MR imaging process. The FWHM of the magnitude PSF does not differentiate between blurring and high-pass filtering effects.

What, then, does the magnitude PSF describe, and can it be used for any characterization of the MRI process? The magnitude PSF has some general value in that it describes the absolute level of influence that neighboring positions in the original pattern have on each other s value in the image. The problem is that it fails to characterize the nature of this influence (e.g. blurring or high-pass filtering), which depends on the signs of the components, the phase and the overall shape of the underlying complex PSF.

### Applicability of our simplified model

#### Approximation of fMRI as a convolution with the real component of the complex imaging PSF

We have shown that the MR imaging process can be approximated by a convolution with the real component of the complex imaging PSF. This approximation works well if the original pattern constitutes a low amplitude spatially varying pattern superimposed on a constant, spatially homogeneous background of a higher amplitude. For response amplitudes of 5%, we found the typical RMSE relative to the standard deviation of the pattern imaged by the complete MRI process to be well below 1%. This scenario holds for fMRI, where gray matter has a relatively uniform baseline intensity and the focus is on superimposed signal changes of approximately 1%-5%. In contrast, this approximation may not be as appropriate for structural MRI where the low signal background, as well as objects of varying size and intensity, are of interest.

### Separation of MRI sampling and signal decay

The separation of MR sampling and signal decay makes it possible to consider their respective effects separately. MR sampling with no decay does not automatically result in a blurred image. If the voxel width is sufficiently small for sampling the larger part of the original spatial frequencies, the MRI sampling will have no blurring effect.

For example, the spatial frequency content of the BOLD response of cortical columns is limited due to the smoothness of the neuronal columnar organizations and the spreads of the neurophysiological and hemodynamic responses. Therefore, the blurring effect of fMRI sampling with adequately small voxels can be neglected. The condition is that the voxels are sufficiently small, such that they can capture the main (peak) frequency of a columnar organization and the frequencies showing elevated power around it.

When untangled from MR sampling, signal decay can be described as a blurring process which we model as Gaussian blurring. This is a simplifying model, with precision that varies with the actual imaging MTF. However, the deviations of this simplifying model relative to the complete fMRI process are small for typical signal changes (fMRI response) and noise levels in fMRI.

Importantly, it results in a simple and intuitive characterization of the blurring associated with signal decay that makes it possible to compare it to previously reported FWHMs of PSFs associated with the entire BOLD fMRI process (BOLD PSF) (Chaimow et al., 2017; Engel et al., 1997; Parkes et al., 2005; Shmuel et al., 2007). It further makes it possible to decompose the PSF of fMRI into two components: one caused by k-space sampling and signal decay, and the other caused by a physiological fMRI contrast-dependent spread.

### Our proposed model applies to both isotropic and anisotropic cortical columns

We have used the example of ocular dominance columns (ODC) in order to show the limitations of using the magnitude PSF for characterizing the effective spatial resolution of fMRI (Figure 2). However, our characterization of the MR imaging process as Gaussian blurring followed by MR sampling with no signal decay does not depend on this specific example. In particular because we only considered the phase-encode dimension, we were able to use one dimensional columnar patterns. These one dimensional patterns can be regarded as general models of columnar patterns, characterized by a main pattern frequency and a level of irregularity.

### Applicability of our proposed model to other fMRI and MRI methods

In addition to GE and SE BOLD fMRI, other methods such as 3D GRASE (Feinberg et al., 2008), VASO (Lu et al., 2003) and ASL (Detre et al., 1992) have been used for high resolution imaging (Duong et al., 2001; Huber et al., 2015; Zimmermann et al., 2011). Do our findings generalize and account for MRI sampling and signal decay in these methods? Our analysis of the imaging PSF depends on two conditions. The first is the shape of signal decay during linear sampling of k-space. This condition holds when using different echo times or preparatory pulses that only affect the absolute magnitude of the magnetization to which the modeled 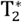 or 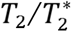 decay is applied. Our derivations are valid for such scenarios. The second is the validity of a linear approximation of the response pattern, which holds if the response is composed of a small amplitude response pattern relative to a larger amplitude spatially homogenous baseline.

In 3D GRASE, multiple refocusing pulses and subsequent EPI acquisitions (partitions) follow a single excitation pulse. Each partition represents a step in 3D k-space in the direction orthogonal to what is commonly considered the slice planes. The signal decay within each partition is proportional to that of a single SE EPI acquisition (Fig. 1, first row, SE). Across partitions, the amplitude changes according to *T*_2_ decay.

Consequently, the entire 3D k-space modulation due to signal decay can be separated into a product of decay between and within partitions. Furthermore, the separability of dimensions in the Fourier transform implies that these respective components determine the spatial filtering due to signal decay across slices (between partition decay) and within slices (within partition decay).

As a result, our SE findings are valid for the effective in-plane resolution of 3D GRASE acquisitions. In order to approximate the effective in-plane spatial resolution of 3D GRASE, one needs to consider the total readout for a single partition and refer to the SE results in Figure 7. However, the effective spatial resolution across slices is determined by a decay similar to GE acquisition, but with a time constant of *T*_2_ instead of 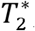, and with a total read-out duration that covers the acquisition of all partitions.

VASO, a method that indirectly measures changes in cerebral blood volume, applies an inversion recovery pulse prior to a normal GE or SE EPI sequence. The effect of the inversion recovery pulse is that all signals from blood are nulled when the excitation pulse occurs. This, in addition to the often used very short echo time, results in a change of scaling but not in a change of the shape of 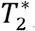 decay (for GE) or 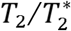 decay (for SE). In addition, VASO signal changes are small, on the order of -1% (Lu and van Zijl, 2012). Together, these features allow us to apply our results directly to VASO imaging. Therefore, Figure 7 provides the effective spatial resolution of VASO acquired by means of GE or SE imaging.

For cerebral blood flow imaging, e.g. using ASL, the situation is different. If EPI acquisition is used, the signal decay follows our analysis, therefore complying with the first condition as described above. However, cerebral blood flow changes are on the order of 20%-60% which does not follow the second condition, making our linear approximation much less accurate.

### How does the effective spatial resolution influence functional imaging?

#### Signal amplitude reductions

In the current study, we focus on signal decay-dependent modulation of amplitudes of spatial frequency components relative to each other. However, signal decay also causes overall amplitude decreases that bring about a reduced signal to thermal noise ratio (SNR). Such amplitude decreases depend on total read-out duration. In addition, read-out duration per read-out line determines receiver bandwidth, with higher bandwidth (shorter read-out duration) resulting in increased noise. Both factors need to be considered in order to find an optimal total read-out duration that maximizes SNR (Qin, 2012), all within the constraints (and potential effects) of matrix size, field of view, echo time, and gradient strength.

### SNR associated with a spatial frequency

SNR and effective spatial resolution can be considered together by evaluating the SNR at a spatial frequency. For detecting or decoding stimulus-specific responses, SNR needs to be high for at least part of the spatial frequencies that contribute to stimulus-specific responses, independent of whether the overall image is blurred. However, if one aims to obtain a precise reconstruction of the response pattern, then both high SNR and an image with an undistorted frequency spectrum are necessary.

In this context, it is of interest to discuss the difference between partial Fourier reconstructions employing conjugate symmetry and simple zero-filling. It may appear that the conjugate symmetry reconstruction is always superior to zero-filling due to its reduced blurring effect. Indeed, this is the case if the aim is to image a pattern precisely. However, the situation is different if we consider the SNR at each spatial frequency. Compared to conjugate symmetry reconstruction, zero-filling reduces the amplitude of high spatial frequencies, as the contribution of their omitted components is set to zero. However, the noise at these spatial frequencies is reduced proportionally. Therefore, the spatial frequency-specific SNR is equal between conjugate symmetry and zero filling reconstructions of partial Fourier acquisition. The spatial frequency specific SNR obtained by full k-space acquisition is higher than those obtained by both conjugate symmetry and zero-filling reconstructions of data obtained by partial Fourier acquisition. This is because full k-space acquisition benefits from averaging of independent noise across the negative and positive k-space parts. The result is an expected increase of spatial frequency-specific SNR by a factor of 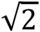 compared to partial Fourier acquisitions, independent of the employed reconstruction method.

### Significance of signal decay blurring relative to the overall BOLD point spread function

The absolute width of the PSF due to signal decay is proportional to voxel width. At ultra-high magnetic field, the in-plane voxel width used for high-resolution fMRI can be as small as 0.5 mm. In Table 1, we compare the FWHM of the resulting PSF due to signal decay to the overall BOLD fMRI PSF as estimated in (Chaimow et al., 2017). Furthermore, we estimated the BOLD PSF while accounting for the effect of signal decay. We considered that consecutive Gaussian convolutions result in a Gaussian convolution with a total width equal to the square root of the sum of squares of individual convolution widths. The results show that the contribution of fMRI signal decay to the overall BOLD fMRI PSF is relatively small. This is especially true, considering that signal decay blurring acts only on the phase-encode direction.

**Table 1.** Comparison of decay dependent imaging PSF and BOLD PSF. This table compares the FWHM of the PSF due to signal decay (fMRI signal decay PSF, as estimated in the current study) to that of the overall BOLD fMRI PSF as estimated in (Chaimow et al., 2017). A voxel width of 0.5 mm is assumed. The bottom row shows the FWHM of the BOLD PSF when accounting for the contribution of fMRI signal decay. These numbers show that the contribution of fMRI signal decay is relatively small. Note that for GE, accounting for the PSF due to signal decay widens the PSF function, due to the high-pas filtering effect of the signal decay in GE BOLD fMRI.

|  | GE | SE |
| --- | --- | --- |
| **Overall BOLD fMRI PSF**<br>(Chaimow et al., 2017) | 0.99 mm | 0.86 mm |
| **fMRI signal decay PSF** | -0.30 mm<br>(high pass filter) | 0.44 mm |
| **BOLD PSF,**<br>**accounting for fMRI signal decay** | 1.03 mm | 0.74 mm |

## Conclusion

We have demonstrated that the FWHM of the absolute values of the complex imaging PSF (magnitude PSF) is a poor and potentially misleading measure for the effect of signal decay on the effective spatial resolution. Instead, we propose to first linearly approximate the typically non-linear process of MR sampling and reconstruction and then to separately consider the effects of two components of the imaging process. The first component is the MR sampling with no signal decay, which acts as a hard low-pass filter. It discards all spatial frequencies higher than the voxel size-dependent maximal sampled spatial frequency and leaves all other frequencies untouched. The second component depends on the signal decay. We have shown that the effect of this second component can be approximated by either Gaussian blurring or high-pass filtering that reverses the effect of a Gaussian blurring. For typical SE parameters at 7 Tesla, we found that the Gaussian blurring attributed to signal decay has a PSF FWHM of 0.89 voxels (0.44 mm for 0.5 mm wide voxels). In contrast, GE at 7 Tesla has a high-pass filter effect, reversing a Gaussian blurring with a PSF FWHM of 0.59 voxels (0.30 mm for 0.5 mm wide voxels). We conclude that signal decay in SE fMRI with full k-space acquisition at 7 Tesla has a more moderate blurring effect compared to the effect implied by the commonly used FWHM of the magnitude PSF. We further conclude that signal decay in GE fMRI at 7 Tesla (and also at other field strengths, not shown) has a high-pass filtering effect, opposite to what can be expected from describing the effect of signal decay on GE fMRI by the FWHM of the corresponding magnitude PSF.

## Acknowledgements

We thank Avery Berman for his comments on an earlier version of the manuscript, and Nicky Tam for English editing. This work was supported by grants from the Natural Sciences and Engineering Research Council of Canada (AS, NSERC Discovery grants RGPIN 375457-09 and RGPIN-2015-05103).

## Appendix A. Modeling and simulating MR imaging

This section provides detailed equations of the MR imaging model as used in our simulations. The theory follows Haacke et al. (1999).

We consider the phase-encode dimension only and analyze it separately from the read-out dimension. This is justified because of the separability of the Fourier transform. Let *y*(*x*) be a spatial pattern and *s*(*k*)=ℱ *y*(*x*) its k-space representation obtained by Fourier transform. Furthermore, let *L* be the field-of-view and *N* = 2*n* the matrix size with voxel width ∆*x* = *L/N*.

### MR sampling with no signal decay

MRI samples k-space in steps of ∆*k* = 1*/L* from -*n*∆*k* to (*n* -1)∆*k*. The reconstructed imaged pattern 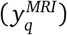, where *q* ∈ [-*n*, …, *n*-1], is obtained by taking the absolute values of an inverse discrete Fourier transform, such that

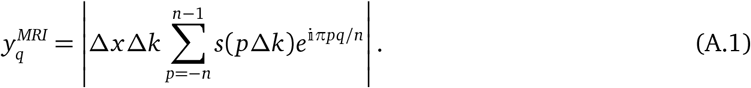

Defining 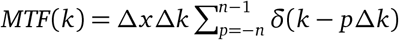, A.1 can be rewritten as:

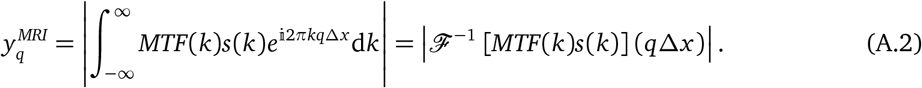

This shows that *MTF*(*k*) is the modulation-transfer function of the linear part of the MRI process (up to taking absolute values). As such it has an associated point-spread function *psf*(*x*)=ℱ^-1^ [*MTF*(*k*)], allowing us to express A.2 as a convolution:

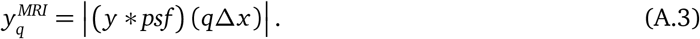

### MR sampling in the presence of signal decay

Let *t*(*k*) be the time that an individual k-space point *k* is being acquired and let *f*(*k*)=*f*(*t*(*k*))be the relative signal decay amplitude at that time. Such an acquisition in the presence of signal decay results in an effective k-space representation *f* (*k*)*s*(*k*), changing A 1 to

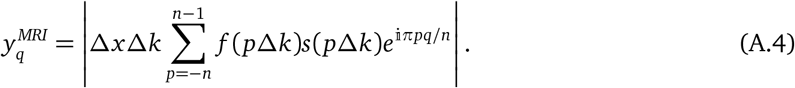

In this situation the MR imaging equations A.2 and A.3 still a_P_pply, if we absorb *f* (*k*) into the modulation-transfer function, now defined as 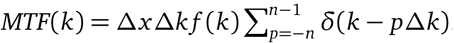.

## Appendix B. Linear approximation of the MR imaging process

Let *r*: [-*L*/2, *L*/2] ⟶ ℝ be a spatial pattern of relative responses of BOLD contrast and let

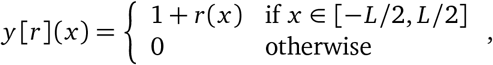

be the associated pattern of absolute BOLD signal.

First we compute the result of convolving such a signal pattern with a point-spread function defined according to Appendix A.

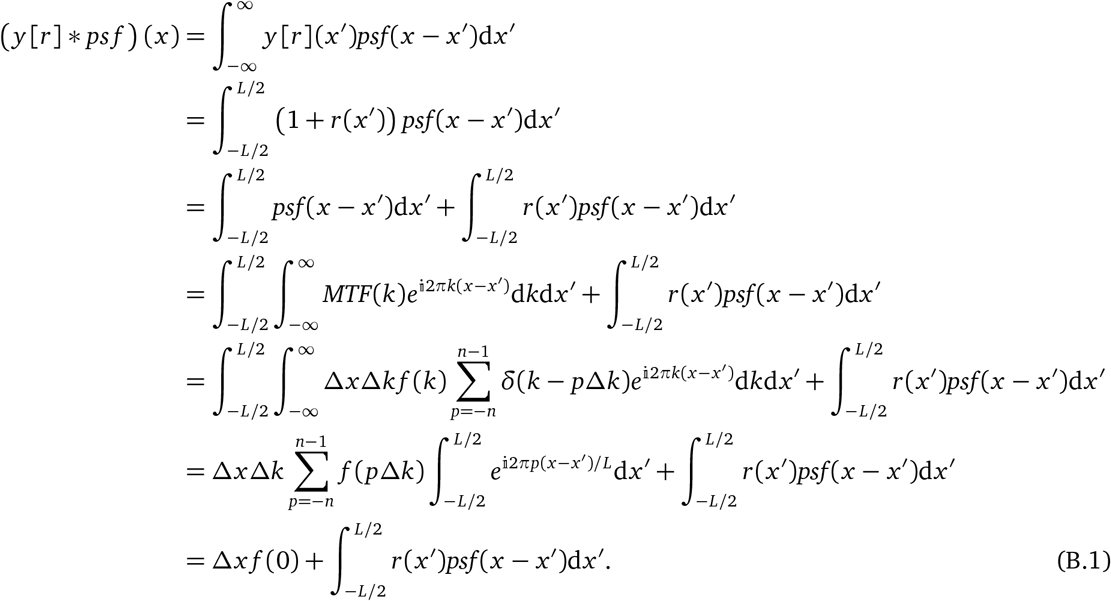

We define an operator *MRI* that models the MRI acquisition process by mapping the spatial pattern *r*(*x*) onto a measured MRI pattern 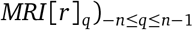 according to A.3, such that

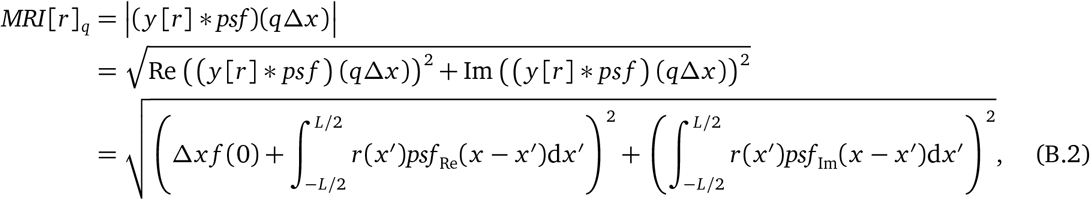

where *psf*_Re_(*x*)=Re *psf* (*x*) and *psf* _Im_(*x*)=Im *psf* (*x*).

We also note that

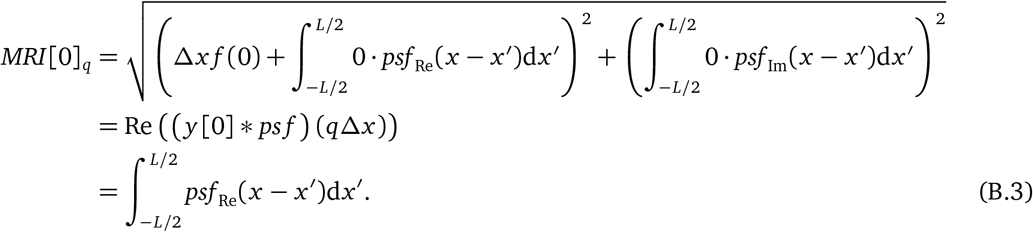

In order to linearly approximate the MR imaging process, we compute *MRI*^′^[0][*r*]_*q*_, the functional derivative of *MRI*[*r*]_*q*_ with respect to *r*, evaluated at *r*_0_ = 0 (representing no response, only baseline). This functional derivative is a linear operator that maps a response pattern *r*(*x*) onto an MRI response pattern (*MRI*^′^[0][*r*]*q*)-*n*≤*q*≤*n*-1, resulting in a linear approximation of the true MRI measurement according to *MRI*[*r*]_*q*_ ≈*MRI*[0]_*q*_ + *MRI*^′^[0][*r*]_*q*_.

We calculate *MRI*^′^[0][*r*]_*q*_ from B.2 using the chain rule and the fact that 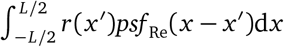 is already a linear operator on *r*:

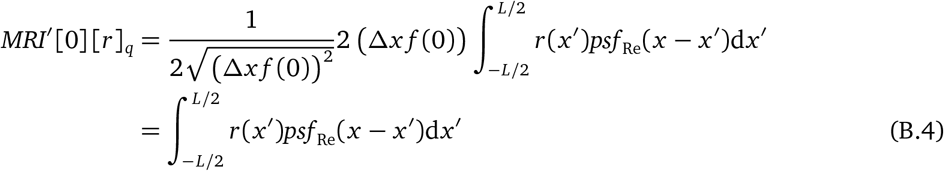

Finally, using B.3 and B.4 the linear approximation of *MRI*[*r*]_*q*_ is

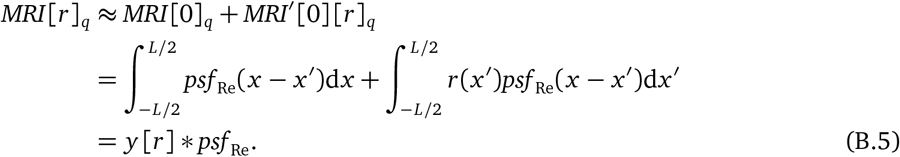

## References

Chaimow, D., Yacoub, E., Uğurbil, K., Shmuel, A., 2017. Spatial specificity of the functional MRI blood oxygenation response relative to neuronal activity. Neuroimage. 10.1016/j.neuroimage.2017.08.077

Chaimow, D., Yacoub, E., Uğurbil, K., Shmuel, A., 2011. Modeling and analysis of mechanisms underlying fMRI-based decoding of information conveyed in cortical columns. Neuroimage 56, 627–642. doi:10.1016/j.neuroimage.2010.09.037

Cheng, K., Waggoner, R.A., Tanaka, K., 2001. Human ocular dominance columns as revealed by high-field functional magnetic resonance imaging. Neuron 32, 359–374.

Constable, R.T., Gore, J.C., 1992. The loss of small objects in variable TE imaging: implications for FSE, RARE, and EPI. Magn Reson Med 28, 9–24.

Detre, J.A., Leigh, J.S., Williams, D.S., Koretsky, A.P., 1992. Perfusion imaging. Magn Reson Med 23, 37–45.

Duong, T.Q., Kim, D.S., Uğurbil, K., Kim, S.-G.G., 2001. Localized cerebral blood flow response at submillimeter columnar resolution. Proceedings of the National Academy of Sciences 98, 10904–10909. doi:10.1073/pnas.191101098

Engel, S.A., Glover, G.H., Wandell, B.A., 1997. Retinotopic organization in human visual cortex and the spatial precision of functional MRI. Cereb Cortex 7, 181–192. doi:10.1093/cercor/7.2.181

Farzaneh, F., Riederer, S.J., Pelc, N.J., 1990. Analysis of T2 limitations and off-resonance effects on spatial resolution and artifacts in echo-planar imaging. Magn Reson Med 14, 123–139.

Feinberg, D.A., Harel, N., Ramanna, S., Uğurbil, K., Yacoub, E., 2008. Sub-millimeter Single-shot 3D GRASE with Inner Volume Selection for T2 weighted fMRI applications at 7 Tesla. Proc. Intl. Soc. Mag. Reson. Med. 16, 1–1.

Goodyear, B.G., Menon, R.S., 2001. Brief visual stimulation allows mapping of ocular dominance in visual cortex using fMRI. Hum Brain Mapp 14, 210–217.

Haacke, M.E., 1987. The effects of finite sampling in spin-echo or field-echo magnetic resonance imaging. Magn Reson Med 4, 407–421.

Haacke, M.E., Brown, R.W., Thompson, M.R., 1999. Magnetic resonance imaging: physical principles and sequence design.

Huber, L., Goense, J., Kennerley, A.J., Trampel, R., Guidi, M., Reimer, E., Ivanov, D., Neef, N., Gauthier, C.J., Turner, R., Möller, H.E., 2015. Cortical lamina-dependent blood volume changes in human brain at 7 T. Neuroimage 107, 23–33. doi:10.1016/j.neuroimage.2014.11.046

Kemper, V.G., De Martino, F., Vu, A.T., Poser, B.A., Feinberg, D.A., Goebel, R., Yacoub, E., 2015. Sub-millimeter T2 weighted fMRI at 7 T: comparison of 3D-GRASE and 2D SE-EPI. Front Neurosci 9, 163. doi:10.3389/fnins.2015.00163

Lu, H., Golay, X., Pekar, J.J., van Zijl, P.C.M., 2003. Functional magnetic resonance imaging based on changes in vascular space occupancy. Magn Reson Med 50, 263–274. doi:10.1002/mrm.10519

Lu, H., van Zijl, P.C.M., 2012. A review of the development of Vascular-Space-Occupancy (VASO) fMRI. Neuroimage 62, 736–742. doi:10.1016/j.neuroimage.2012.01.013

Menon, R.S., Ogawa, S., Strupp, J., Uğurbil, K., 1997. Ocular dominance in human V1 demonstrated by functional magnetic resonance imaging. J Neurophysiol 77, 2780–2787.

Nasr, S., Polimeni, J.R., Tootell, R.B., 2016. Interdigitated Color-and Disparity-Selective Columns within Human Visual Cortical Areas V2 and V3. J Neurosci. 36:1841–57.

Oshio, K., Singh, M., 1989. A computer simulation of T2 decay effects in echo planar imaging. Magn Reson Med 11, 389–397.

Parkes, L.M., Schwarzbach, J.V., Bouts, A.A., Deckers, R.H.R., Pullens, P., Kerskens, C.M., Norris, D.G., 2005. Quantifying the spatial resolution of the gradient echo and spin echo BOLD response at 3 Tesla. Magn Reson Med 54, 1465–1472. doi:10.1002/mrm.20712

Polimeni, J.R., Fischl, B., Greve, D.N., Wald, L.L., 2010. Laminar analysis of 7T BOLD using an imposed spatial activation pattern in human V1. Neuroimage 52, 1334–1346. doi:10.1016/j.neuroimage.2010.05.005

Qin, Q., 2012. Point spread functions of the T2 decay in k-space trajectories with long echo train. Magn Reson Imaging 30, 1134–1142. doi:10.1016/j.mri.2012.04.017

Rojer, A., Schwartz, E., 1990. Cat and monkey cortical columnar patterns modeled by bandpass-filtered 2D white noise. Biol Cybern 62, 381–391.

Shmuel, A., Chaimow, D., Raddatz, G., Uğurbil, K., Yacoub, E., 2010. Mechanisms underlying decoding at 7 T: ocular dominance columns, broad structures, and macroscopic blood vessels in V1 convey information on the stimulated eye. Neuroimage 49, 1957–1964. doi:10.1016/j.neuroimage.2009.08.040

Shmuel, A., Yacoub, E., Chaimow, D., Logothetis, N.K., Uğurbil, K., 2007. Spatiotemporal point-spread function of fMRI signal in human gray matter at 7 Tesla. Neuroimage 35, 539–552. doi:10.1016/j.neuroimage.2006.12.030

Uludağ, K., Müller-Bierl, B., Uğurbil, K. 2009. An integrative model for neuronal activity-induced signal changes for gradient and spin echo functional imaging. Neuroimage 48, 150–165. doi:10.1016/j.neuroimage.2009.05.051

Yacoub, E., Duong, T.Q., van de Moortele, P.F., Lindquist, M., Adriany, G., Kim, S.-G.G., Uğurbil, K., Hu, X.P., 2003. Spin-echo fMRI in humans using high spatial resolutions and high magnetic fields. Magn Reson Med 49, 655–664. doi:10.1002/mrm.10433

Yacoub, E., Harel, N., Uğurbil, K., 2008. High-field fMRI unveils orientation columns in humans. Proceedings of the National Academy of Sciences 105, 10607–10612. doi:10.1073/pnas.0804110105

Yacoub, E., Shmuel, A., Logothetis, N.K., Uğurbil, K., 2007. Robust detection of ocular dominance columns in humans using Hahn Spin Echo BOLD functional MRI at 7 Tesla. Neuroimage 37, 1161–1177.

Yacoub, E., Shmuel, A., Pfeuffer, J., van de Moortele, P.F., Adriany, G., Andersen, P., Vaughan, J.T., Merkle, H., Uğurbil, K., Hu, X. 2001. Imaging brain function in humans at 7 Tesla. Magn Reson Med 45, 588–594. doi:10.1002/mrm.1080

Zimmermann, J., Goebel, R., De Martino, F., van de Moortele, P.-F., Feinberg, D., Adriany, G., Chaimow, D., Shmuel, A., Uğurbil, K., Yacoub, E., 2011. Mapping the organization of axis of motion selective features in human area MT using high-field fMRI. PLoS ONE 6, e28716. doi:10.1371/journal.pone.0028716

